# Ubiquitinated histone H2B as gatekeeper of the nucleosome acidic patch

**DOI:** 10.1101/2024.02.22.581437

**Authors:** Chad W. Hicks, Sanim Rahman, Susan L. Gloor, James K. Fields, Natalia Ledo Husby, Anup Vaidya, Keith E. Maier, Michael Morgan, Michael-Christopher Keogh, Cynthia Wolberger

**Affiliations:** Department of Biophysics & Biophysical Chemistry, Johns Hopkins University School of Medicine, 725 N. Wolfe Street, Baltimore, MD 21205, United States; EpiCypher Inc., 6 Davis Drive, Suite 755, Durham, NC 27709, United States

**Author notes:** Michael Morgan, Stablix Inc., 201 Brookline Avenue, Suite 1, Boston, MA, 02215, United States.

## Abstract

Monoubiquitination of histones H2B-K120 (H2BK120ub) and H2A-K119 (H2AK119ub) play opposing roles in regulating transcription and chromatin compaction. H2BK120ub is a hallmark of actively transcribed euchromatin, while H2AK119ub is highly enriched in transcriptionally repressed heterochromatin. Whereas H2BK120ub is known to stimulate the binding or activity of various chromatin-modifying enzymes, this post-translational modification (PTM) also interferes with the binding of several proteins to the nucleosome H2A/H2B acidic patch via an unknown mechanism. Here we report cryoEM structures of an H2BK120ub nucleosome showing that ubiquitin adopts discrete positions that occlude the acidic patch. Molecular dynamics simulations show that ubiquitin remains stably positioned over this nucleosome region. By contrast, our cryoEM structures of H2AK119ub nucleosomes show ubiquitin adopting discrete positions that minimally occlude the acidic patch. Consistent with these observations, H2BK120ub, but not H2AK119ub, abrogates nucleosome interactions with acidic patch-binding proteins RCC1 and LANA, and single-domain antibodies specific to this region. Our results suggest a mechanism by which H2BK120ub serves as a gatekeeper to the acidic patch and point to distinct roles for histone H2AK119 and H2BK120 ubiquitination in regulating protein binding to nucleosomes.

## INTRODUCTION

The nucleosome is the fundamental organizational unit of chromatin and comprises two copies each of histones H2A, H2B, H3, and H4 wrapped by ∼147 base pairs (bp) of DNA. Nuclear proteins interact with the nucleosome at various sites including the histone core, histone tails, nucleosomal DNA, and extra-nucleosomal linker DNA (1,2). The core histones are subject to a wide array of post-translational modifications (PTMs) that play a central role in regulating transcription (3), DNA replication (4), chromatin compaction (5), and the DNA damage response (6). PTMs ranging from small chemical groups, such as methyl, acetyl, and phosphoryl, to the attachment of 76-amino acid ubiquitin (7), modulate protein binding to chromatin and impact higher-order chromatin structure (8,9), thereby regulating downstream biological processes (10,11).

The nucleosome contains a conserved cluster of negatively charged residues in histones H2A and H2B, known as the nucleosome acidic patch (12), that is a hotspot for interactions with proteins that regulate transcription, the cell cycle, and the DNA damage response (reviewed in (2)). Structural studies have shown that proteins primarily engage the nucleosome acidic patch with arginine residues (2), as first seen in the crystal structure of the Ran GTPase, RCC1, bound to a nucleosome (13). A recent proteomics study identified many additional proteins that bind the acidic patch and are involved in epigenetic regulation, transcription, and cell-cycle regulation (14).

Monoubiquitination of histones H2B-K120 (H2BK120ub) and H2A-K119 (H2AK119ub) play opposing roles in transcription regulation and chromatin compaction (15,16). H2BK120ub is enriched in actively transcribed genes (17,18), where it stimulates methylation of histone H3-K4 (17,19,20) and H3-K79 (18,21,22). H2BK120ub is dynamically regulated throughout the cell cycle (23), serving as a signal for the DNA damage response (24) and chromatin decompaction (8). H2AK119ub is enriched at heterochromatic regions (25), where it stimulates trimethylation of H3K27 (26–28), a repressive mark (29), and promotes chromatin compaction (30,31).

There have been several reports that H2BK120ub can interfere with the binding of proteins to nucleosomes, including human RCC1, yeast Sir3, and Kaposi’s sarcoma-associated herpesvirus latency-associated nuclear antigen (LANA) peptide (32–34). All three proteins contact the nucleosome acidic patch, which raises the possibility that the ubiquitin conjugated to H2BK120 interferes with their binding to this surface (13,35,36). However, a role for H2BK120ub in blocking protein binding seems at odds with the presumed mobility of ubiquitin, given the flexibility of its carboxy-terminal tail and of the aliphatic lysine side-chain to which it is covalently linked (37). Such flexibility could, in principle, allow ubiquitin to adopt positions on the nucleosome surface that would not clash with protein binding.

To gain insight to the structural basis by which H2BK120ub could interfere with protein binding to the nucleosome, we determined cryoEM structures of nucleosomes containing ubiquitin conjugated to H2B-K120. We found that the ubiquitin adopts several distinct positions that partially occlude the nucleosome acidic patch. Molecular dynamics (MD) simulations indicate that the ubiquitin conjugated to H2B-K120 remains stably associated over the nucleosome acidic patch, in positions that would be expected to interfere with protein binding. We also determined cryoEM structures of H2AK119ub nucleosomes and found that, while the ubiquitin adopts two distinct positions, neither significantly occludes the acidic patch. In vitro studies showed that H2BK120ub, but not H2AK119ub, dramatically reduces nucleosome binding by RCC1, LANA peptide, and single domain antibodies that specifically contact the acidic patch. Our results establish a mechanism by which H2BK120ub can regulate protein binding to the nucleosome acidic patch, thereby impacting a major hub of chromatin interactions.

## MATERIAL AND METHODS

### Expression and purification of ubiquitin

Ubiquitin (G76C) was expressed and purified as previously described (38).

### Expression and purification of histones

Expression plasmids for *Xenopus laevis* histones H2A, H2B, H3 and H4 were a gift from Gregory Bowman (Johns Hopkins University). Histones were expressed and purified as previously described (39), with the following modifications. Thawed cell pellet was lysed with multiple passes through a Microfluidizer (Microfluidics). After lysis, centrifugation, and multiple washes using a Triton X-100 containing buffer, washed cell pellets were resuspended in a buffer containing 20 mM HEPES pH 7.5, 7 M Guanidine HCl, 10 mM dithiothreitol (DTT). Resuspended washed cell pellets were size exclusion purified using a buffer containing 10 mM Tris pH 8.0, 7 M urea, 1 mM EDTA, 5 mM b-mercaptoethanol (BME). Fractions were pooled and injected onto a 2-column series with a HiTrap Q-XL column connected upstream of a HiTrap SP-XL column using a buffer containing 20 mM Tris pH 7.8, 7 M urea, 1 mM EDTA, 5 mM BME). The Q-XL column was removed from the system, and then the histones were eluted from the SP-XL column with a gradient of 0 – 1 M NaCl on an ÄKTA Pure (Cytiva) chromatography system.

### Preparation of ubiquitinated histone H2BK120 and H2AK119

Dichloroacetone (DCA)-linked H2BK120ub and H2AK119ub were prepared as previously described (38).

### Preparation and purification of dimethylated histone H3-K79 (H3K79me2)

A methyl-lysine analog of H3K79me2 (H3K_C_79me2) was prepared by alkylating the cysteine in histone H3K79C as previously described (40) and lyophilized. To purify the reaction product, the lyophilized sample was dissolved in filtered 7 M guanidine HCl and loaded onto a PROTO 300 C4 reverse-phase column (Higgins Analytical) with a buffer containing 0.1% TFA and eluted with a gradient of 0-90% acetonitrile on an HPLC (Waters). The peaks corresponding to H3K_C_79me2 and unreacted histone H3-K79C were collected and identified. Identity and purity were determined by digesting a sample with trypsin and analyzing its mass/charge ratio using Matrix Assisted Laser Desorption/Ionization - Mass Spectrometry (MALDI-MS). The H3K_C_79me2 sample was dialyzed into 5 mM BME, lyophilized, and stored at -20°C.

### Purification of Widom 601 DNA

The Widom 601 DNA sequence (145 bp) (41) was expressed in *E. coli* XL-1 Blue using the pST55-16x601 plasmid (13). The 601 sequence was expressed, purified, and isolated as described (42). The 601 sequence used is:

5’−ATCGATGTATATATCTGACACGTGCCTGGAGACTAGGGAGTAATCCCCTTGGCGGTTAAAACGCGGGGGACAGCGCGTACGTGCGTTTAAGCGGTGCTAGAGCTGTCTACGACCAATTGAGCGGCCTCGGCACCGGGATTCTGAT−3’.

601 DNA flanked by 20 bp of linker DNA (185 bp; 20-N-20) was amplified using polymerase chain reaction (PCR) using primers (IDT DNA) containing 20 bp overhangs. The primers used were:

Forward:

5’−biosg−GTCGCTGTTCGCGACCGGCAATCGATGTATATATCTGACACGTGCC−3’ Reverse: 5’−GACCCTATACGCGGCCGCCCATCAGAATCCCGGTGCCGAG−3’

Phusion polymerase was used to amplify 100 μL reaction volumes with standard PCR parameters. The PCR product was precipitated by combining 100% EtOH, PCR mix, and 3 M sodium acetate pH 5.2 (10:1:20) at -80°C for 1 hour. Precipitated DNA was collected by centrifugation and the supernatant was removed. The pellet was washed with 70% EtOH and air dried before resuspending in TE Buffer (10 mM Tris pH 8.0, 1 mM EDTA). The PCR product was purified with PCI (phenol:chloroform:isoamyl alcohol 25:24:1). PCI was added to an equal volume of resuspended PCR product pellet, vortexed, and centrifuged to separate the phases. The organic phase was removed and the aqueous phase was extracted twice more the same way. All of the removed organic phases were pooled, back-extracted with TE buffer to collect any PCR product left behind in the organic phase, and all the aqueous phases were combined and saved as the purified PCR product. Purified PCR product, 3 M sodium acetate pH 5.2, and 100% EtOH was combined in a 10:1:20 ratio and placed at -20°C overnight. The precipitated purified PCR product was then centrifuged, the pellet washed with 70% EtOH, centrifuged again, and allowed to dry. This pellet of purified 185bp 601 DNA was then resuspended in MilliQ (Sigma) water and stored at -20°C.

### Preparation of nucleosomes

Nucleosomes for structural studies (unmod-Nuc 145bp/185bp, H2AK119ub Nuc 145/185bp, H2BK120ub Nuc 145/185bp, H2BK120ub+H3K_C_79me2 Nuc 185bp) were reconstituted as previously described (39), with the modifications described below.

For each sample, histone octamer and Widom 601 DNA was combined in an octamer:DNA molar ratio of 1.2:1 in a buffer containing 10 mM Tris pH 7.5, 2 M KCl, 1 mM EDTA, 1 mM DTT, such that final DNA concentration was 6 μM. Nucleosomes were then assembled by salt gradient dialysis, reducing the salt concentration to 0.25 M KCl over 24 hours.

Precipitate was removed by centrifugation and nucleosome purity assessed by native gel electrophoresis. Nucleosome samples that showed excess free DNA or higher-order species were further purified by loading onto a SK DEAE-5PW column (TOSOH biosciences) equilibrated in buffer containing 10 mM Tris pH 7.5, 0.25 M KCl, 0.5 mM EDTA, 1 mM DTT, and eluted with a gradient of 0.25 – 0.6 M KCl on an Agilent HPLC instrument. Purified nucleosome was dialyzed into a buffer containing 20 mM HEPES pH 7.5, 25 mM KCl, 1 mM EDTA, 1 mM DTT, 20% glycerol, flash frozen in liquid nitrogen, and stored at -80°C.

Nucleosomes for dCypher™ Luminex nucleosome binding assays (unmodified (*EpiCypher* 16-0006); acidic patch mutant H2A(E61A) (*EpiCypher* 16-0029); H2AK119ub1 (*EpiCypher* 16-0395); or H2BK120ub1 (*EpiCypher* 16-0396)) were generated on 5’ biotinylated DNA (147bp of 601 nucleosome positioning sequence) as previously described (43–45) and confirmed by SDS-PAGE, immunoblotting and mass spectrometry. The H2B-K120 and H2A-K119 ubiquitinated histones used for these binding studies possess a native gamma-lysine isopeptide linkage (44,46).

### Purification of RCC1

A plasmid encoding RCC1 fused to an N-terminal hexahistidine (6xHis) tag was transformed into *E. coli* BL21(DE3)Rosetta2-pLysS cells. The colonies were used to inoculate 5 mL volumes of media, which were expanded to 1 L volumes for full-scale growth. Cultures were grown at 37°C and 200 RPM in 2x Yeast Extract Tryptone (2XYT) media supplemented with ampicillin and chloramphenicol. Cultures were induced by addition of 1 mM isopropyl-ß-D-thiogalactopyranoside (IPTG) when they reached an OD_600_ of 0.4 – 0.6, and were grown for an additional 18 hours at 18°C. Cells were harvested by centrifugation, resuspended in buffer containing 20 mM HEPES pH 7.5, 300 mM NaCl, 50 mM imidazole, 10% glycerol, 1 mM DTT, 1 tablet/50 mL Complete Protease Inhibitor Cocktail (Millipore Sigma #11836153001), flash-frozen in liquid nitrogen, and stored at -80°C.

The frozen cell pellet suspension was thawed in a water bath and an equal volume of buffer containing 20 mM HEPES, pH 7.5, 300 mM NaCl, 50 mM imidazole, 1 mM DTT and 0.2 mM PMSF added. Diluted cell suspension was lysed by sonication for three rounds each of 1 min total processing time (5sec on/ 10 sec off) at 40% power. The resulting whole cell extract was centrifuged at 17,000 RPM and the supernatant filtered using a 1.1 μm filter. The resulting clarified extract was loaded onto a 5 mL HisTrap HP (Cytiva) column equilibrated in 20 mM HEPES, pH 7.5, 300 mM NaCl, 50 mM imidazole, 1 mM DTT, and eluted with a gradient of 0.05 - 1 M imidazole on an ÄKTA Pure instrument (Cytiva). Eluted protein was diluted with 20 mM HEPES pH 7.5, 10 % glycerol, 1 mM DTT, 0.2 mM PMSF to a final salt concentration of 50 mM NaCl and filtered using a 1.1 μm filter. The resulting protein was loaded onto 5 mL HiTrap SP-HP (Cytiva) cation exchange column equilibrated in 20 mM HEPES pH 7.5, 50 mM NaCl, 10% glycerol, 1 mM DTT, 0.2 mM PMSF and eluted with a gradient of 0.05 - 2 M NaCl. Eluted protein was dialyzed overnight in 20 mM HEPES, 150 mM NaCl, 10% glycerol, 1 mM DTT and 0.2 mM PMSF, concentrated, flash-frozen in liquid nitrogen, and stored at -80°C.

### CryoEM sample preparation

Nucleosome containing H2BK120ub and 145 bp Widom 601 DNA was thawed on ice and buffer exchanged to nucleosome storage buffer (20 mM HEPES pH 7.8, 50 mM NaCl, 1 mM DTT) using a Zeba Spin Desalting Column (Thermo #89882). Quantifoil R 2/2 copper 200 mesh grids (Electron Microscopy Sciences #Q2100CR2) were glow-discharged for 45 seconds at 15 mA using a PELCO easiGLOW Glow Discharge System to apply a negative charge to their surface. Then 3 μL of sample at 0.52 mg/mL was applied to the grid, immediately blotted for 3.5 seconds with a blot force of 5, and plunge-frozen in liquid ethane using a Vitrobot Mark IV (Thermo Fisher) set at 100% humidity and 4°C.

Nucleosome containing histone H2BK120ub, dimethylated histone H3K79 analog (H3K_C_79me2) and the 145 bp Widom 601 DNA with 20 bp linkers (185 bp) was thawed on ice and buffer exchanged into nucleosome storage buffer. Quantifoil R 2/2 copper 200 mesh grids were glow-discharged for 30 seconds at 15 mA, after which 3 μL of sample at 1.50 mg/mL was applied to the grid, left to adsorb for 90 seconds, then blotted for 3 seconds with a blot force of 5, and plunge-frozen in liquid ethane using a Vitrobot Mark IV apparatus set to 100% humidity and 4°C.

Nucleosome containing histone H2AK119ub and the 145 bp Widom 601 DNA was buffer exchanged into nucleosome storage buffer. Quantifoil R 2/2 copper 200 mesh grids were glow-discharged for 45 seconds at 15 mA, after which 3 μL of sample at 0.82 mg/mL was applied to the grid, immediately blotted for 3.5 seconds with a blot force of 5, and plunge-frozen in liquid ethane using a Vitrobot Mark IV apparatus set to 100% humidity and 4°C.

### CryoEM data collection

Data on H2BK120ub nucleosomes were collected at the Beckman Center for Cryo-EM at the Johns Hopkins University School of Medicine using a Thermo Fisher Titan Krios 300 kV electron microscope equipped with a Falcon 4 direct electron detector and a Selectris energy filter. A dataset of 8,392 exposures was collected in counting mode and recorded in Electron Event Representation (EER) format using a magnification of 130kx, pixel size of 0.97 Å, nominal dose of 40 e^-^/ Å^2^, dose rate of 6.42 e^-^/px/s, a defocus range of -0.5 to -2.5 μm, and an energy filter slit width of 10 eV. A multi-shot imaging strategy was used to collect 8 shots per hole, utilizing beam image shift to move between each target.

Data on H2BK120ub+H3K_C_79me2 nucleosomes were collected at the National Cryo-EM Facility (NCEF) at the Frederick National Laboratory for Cancer Research, using a Thermo Fisher Titan Krios 300 kV electron microscope equipped with a Gatan K3 camera and an energy filter. A dataset of 7,782 exposures was collected in super-resolution mode using a magnification of 105kx, pixel size of 0.436 Å, nominal dose of 50 e^-^/ Å^2^, dose rate of 12.33 e^-^/px/s, 40 frames per exposure, a defocus range of -0.75 to -1.75 μm, and an energy filter slit width of 20 eV. A multi-shot imaging strategy was used to collect 3 shots per hole, utilizing beam image shift to move between each target.

Data on H2AK119ub nucleosomes were collected at the Beckman Center for Cryo-EM at Johns Hopkins University School of Medicine using a Titan Krios 300 kV electron microscope equipped with a Falcon 4 direct electron detector and Selectris energy Filter. Two datasets were collected and combined. The first dataset of 5,479 exposures was collected in counting mode and recorded in Electron Event Representation (EER) format using a magnification of 130kx, pixel size of 0.97 Å, nominal dose of 40 e^-^/ Å^2^, dose rate of 6.42 e^-^/px/s, a defocus range of -0.2 to -5.0 μm, and an energy filter slit width of 10 eV. A multi-shot imaging strategy was used to collect 9 shots per hole, utilizing beam image shift to move between each target. A second dataset of 6,732 exposures was collected in counting mode and recorded in EER format using a magnification of 130kx, pixel size of 0.97 Å, nominal dose of 40 e^-^/ Å^2^, dose rate of 6.46 e^-^/px/s, a defocus range of -0.5 to -3.0 μm, and an energy filter slit width of 10 eV. A multi-shot imaging strategy was used to collect 9 shots per hole, utilizing beam image shift to move between each target.

### CryoEM data processing

Data on H2BK120ub nucleosomes were processed in cryoSPARC v4.1 (47). Exposures were motion-corrected and cropped to one-half of their original resolution using Patch Motion Correction. The contrast transfer function (CTF) correction was performed using Patch Motion Correction. Poor quality micrographs were removed using Manually Curate Exposures, yielding 8,149 high-quality micrographs. An initial round of particle picking was performed using Blob Picker and Inspect Picks, then extracted using Extract from Micrographs. A set of 2D templates was created by first performing 2D classification on the extracted particles, then selecting representative views using Select 2D classes. These templates were used to perform a second round of particle picking using Template Picker and Inspect Picks, then extracted using Extract from Micrographs to yield an uncleaned particle stack of 754,004 particles. One round of 2D Classification and Select 2D Classes was performed to remove junk particles by discarding distinctly poor quality 2D classes to yield a partially cleaned particle stack of 550,718 particles. Additional particle cleaning was performed using four parallel 4-structure Ab-Initio Reconstruction jobs, keeping particles from the good classes, which yielded a cleaned particle stack of 329,531 particles. These particles were aligned along their C2 symmetry axis using the Non-Uniform Refinement job (48) and then symmetry-expanded to double the effective number of particles to 659,062 particles. One round of focused 3D Classification was performed with a spherical focus mask centered on the area of blurred ubiquitin density to yield four distinct ubiquitin positions. Individual particle CTF was refined with Local CTF Refinement, image group CTF was refined with Global CTF Refinement, and structures were refined with local refinement to produce H2BK120 ubiquitin position 1 (76,770 particles, 3.32 Å resolution), position 2 (71,307 particles, 3.33 Å resolution), position 3 (73,431 particles, 3.34 Å resolution), and position 4 (65,167 particles, 3.36 Å resolution), The final maps were sharpened with Local Filtering.

Data on H2BK120ub+H3K_C_79me2 nucleosomes were processed in cryoSPARC v3.3 (47). Exposures were motion-corrected and cropped to one-half of their original resolution using Patch Motion Correction. The CTF correction was performed using Patch CTF Estimation. Poor quality micrographs were removed using Manually Curate Exposures, yielding 7,324 high-quality micrographs. An initial round of particle picking was performed using Blob Picker and Inspect Picks, then extracted using Extract from Micrographs. A set of 2D templates was created by first performing 2D Classification on the extracted particles then selecting representative views using Select 2D classes. These templates were used to perform a second round of particle picking using Template Picker and Inspect Picks, then extracted using Extract from Micrographs to yield an uncleaned particle stack of 2,288,109 particles. Two rounds of 2D Classification and Select 2D Classes was performed to remove junk particles by discarding distinctly poor quality 2D classes, yielding a partially cleaned particle stack of 1,034,537 particles. Additional particle cleaning was performed using multiple iterations of Ab-Initio Reconstruction jobs, yielding a cleaned particle stack of 628,403 particles. The particles were aligned along their C2 symmetry axis using the Non-Uniform Refinement job (48) and then symmetry-expanded to double the effective number of particles to 1,256,806 particles. Three rounds of focused 3D Classification were performed with a large spherical focus mask centered on H2BK120 on one face of the nucleosome to yield two distinct ubiquitin positions. Individual particle CTF was refined with Local CTF refinement, and structures were refined with Local Refinement to yield maps showing H2BK120 ubiquitin positions 5 (818,874 particles, 2.93 Å resolution) and 6 (282,239 particles, 3.06 Å resolution). The final maps were sharpened with Local Filtering.

Data on H2AK119ub nucleosomes were processed in cryoSPARC v3.3 (47). Exposures from both datasets were imported with an EER upsampling factor of 2 and motion corrected using Patch Motion Correction. The CTF correction was performed using Patch Motion Correction. Poor quality micrographs were removed using Manually Curate Exposures, yielding 10,018 high-quality micrographs. An initial round of particle picking was performed using Blob Picker and Inspect Picks, then extracted using Extract from Micrographs. A set of 2D templates was created by first performing 2D Classification on the extracted particles then selecting representative views using Select 2D Classes. These templates were used to perform a second round of particle picking using Template Picker and Inspect Picks, then extracted using Extract from Micrographs to yield an uncleaned particle stack of 1,824,939 particles. One round of 2D Classification and Select 2D Classes was performed to remove junk particles by discarding distinctly poor quality 2D classes to yield a cleaned particle stack of 895,614 particles. We note that iterative Ab-Initio Reconstruction jobs for particle cleaning did not improve resolution, so were therefore not used for additional particle cleaning. The particles were aligned along their C2 symmetry axis using the Non-Uniform Refinement job (48) and then symmetry-expanded, doubling the effective number of particles to 1,791,228 particles. These particles were used in a local refinement 3D reconstruction to produce a structure with poor density for ubiquitin, showing a blur of lower resolution density across the surface of the nucleosome. A round of focused 3D classification was performed with a cylindrical focus mask centered on the area of blurred ubiquitin density to yield two distinct ubiquitin positions. An additional round of focused 3D classification was performed for each ubiquitin position using a spherical focus mask centered on each ubiquitin position. Individual particle CTF was refined with Local CTF Refinement, image group CTF was refined with Global CTF Refinement, and structures were refined with local refinement to produce H2AK119 ubiquitin position 1 (102,259 particles, 3.41 Å resolution) and position 2 (68,407 particles, 3.47 Å resolution). The final maps were sharpened with Local Filtering.

### Model building and refinement

Initial models of nucleosomes containing H2BK120ub, H2BK120ub+H3K_C_79me2 and H2AK119ub were constructed by rigid-body fitting models for an unmodified nucleosome (PDB: 4ZUX) and ubiquitin (PDB: 1UBQ) into density maps using ChimeraX (49), and refined using all-atom flexible refinement with strong restraints in *Coot* 0.9.6 (50). Since the particles were subjected to C2 symmetry expansion and 3D classified according to the position of only one of the ubiquitin molecules, the ubiquitin was modeled onto only one side of the nucleosome. Histone tails were extended where density was visible. C-terminal tail residues of ubiquitin were omitted where density was not visible. The modeled ubiquitin includes residues 1-72 in H2Bub position 4, 1-74 in H2Bub position 5, 1-73 in H2Bub position 6, 1-73 in H2Aub positions 1 and 2. The DNA of H2BK120ub+H3K_C_79me2 nucleosomes was extended from 145 bp to 157 bp where density was visible to account for the for the extra-nucleosomal linker DNA present in this sample. All models were further refined in PHENIX (51) using phenix.real_space_refine (52) and validated using the CryoEM Comprehensive Validation module in PHENIX running MolProbity (53). Figures were generated with ChimeraX (49).

### Molecular dynamics simulations

The H2B-ubiquitinated nucleosome system was built using the full-length crystal structure of the *X. laevis* nucleosome (1KX5) (54) and the crystal structure of ubiquitin (1UBQ) (55). Following the approach of Carroll et al. (56), the backbone of the ubiquitin-linked K48 in chain B of PDB entry 3NS8 (57) was aligned with the lysine backbone of histone H2BK120. Next, the sidechain of H2BK120 was rebuilt to have the same geometry as K48 in 3NS8. The ubiquitin C-terminal glycine in chain A of 3NS8 (post-alignment) was then used as the C-terminal glycine of 1UBQ. The unstructured C-terminus of 1UBQ (residues 71-75) was refined using the Modloop server (58) to connect and refine the C-terminal glycine to the rest of 1UBQ. The covalent bond between H2BK120 and the C-terminal glycine of ubiquitin was built using the LEaP program in AMBER20 (59), resulting in a bond between the neutralized lysine sidechain nitrogen and the glycine carbonyl carbon. The structure was solvated in a truncated octahedron box with periodic boundary conditions set to 1.25 nm away from the ubiquitinated nucleosome. Sodium and chloride ions were then added to neutralize the system charge and to reach a final concentration of 150 mM.

All simulations were performed using the AMBER ff14SB (60) force field, including parmbsc1 DNA and CUFIX ion parameter corrections (61,62). These force field parameters have been shown to effectively capture nucleosome dynamics in all-atom MD simulations (63). All water molecules were described using a TIP3P water model. For the peptide bond between H2BK120 and the C-terminal glycine of ubiquitin, we employed the force field parameters parameterized by Carroll et al (56).

Simulations were carried out using the CPU version of Particle-Mesh Ewald Molecular Dynamics (PMEMD) in the AMBER20 software (59). All systems were energy-minimized using a steepest descent gradient method for 10,000 steps with a 500 kJ mol^-1^ nm^-2^ positional restraint applied to all heavy atoms. Systems were then equilibrated as described by Armeev et al. (63): i) 100 ps with positional restraints of 500 kJ mol^−1^ nm^−2^ with 0.5 fs time step; ii) 200 ps with positional restraints of 50 kJ mol^−1^ nm^−2^ with 2 fs time step (and further the same); iii) 200 ps with positional restraints of 5 kJ mol^−1^ nm^−2^; and iv) 200 ps with positional restraints of 0.5 kJ mol^−1^ nm^−2^; v) 200 ps of unrestrained simulations. Systems were equilibrated in the canonical ensemble (NVT), at 300 K and 1 bar using the Langevin thermostat and Parrinello-Rahman barostat. Systems were then simulated with no positional restraints for 1 ns in an isobaric-isothermal (NPT) ensemble at 300 K and 1 bar, using a Langevin thermostat and Monte Carlo barostat (same for production run simulations). All simulations used a 4 fs time step using hydrogen mass repartitioning. Hydrogen bond lengths were constrained using the SHAKE algorithm. The cutoff for non-bonded interactions were set to 12.0 Å, and long-range electrostatic interactions were computed using the particle mesh EWALD (PME) method. Trajectory frames were written every 50 ps. The production run was simulated in duplicate for 400 ns, producing a total of 800 ns of simulation time.

The AmberTools CPPTRAJ package (64) was used to center trajectories. Simulations were visualized using Visual Molecular Dynamics (VMD) (65) and PyMOL. Distance calculations were performed using the MDAnalysis package (66). To calculate the distance between ubiquitin and the nucleosome acidic patch, the distance between the center of mass of each group was used. The center of mass of ubiquitin was based on the backbone atoms of regions with secondary structure. The acidic patch center of mass was based on the backbone heavy atoms of the following residues of histone H2A: E61, E64, D90, E91, and E92. The ubiquitin molecules on both faces of the nucleosome were used from each trajectory. The distances sampled between ubiquitin and the acidic patch are presented as a probability distribution function using the seaborn package in python. To generate ubiquitin clusters, we used the K-means clustering algorithm available in the AmberTools CPPTRAJ package (64). Using both trajectory replicates, we generated 10 clusters based on the RMSD of the backbone heavy atoms of ubiquitin containing secondary structure. Since there are two ubiquitin molecules on the nucleosome, the ubiquitin molecule that had the highest RMSD was used to generate the RMSD clusters. For the calculation of ubiquitin RMSD, all trajectory frames were aligned to the backbone heavy atoms of the histone octamer core in the cryoEM structure of the ubiquitinated nucleosome.

### Electrophoretic mobility shift assays

Binding reactions were prepared in 12 μL volumes by combining RCC1 and nucleosome in binding buffer (20 mM HEPES pH 7.6, 50 mM NaCl, 5% sucrose, 1 mM DTT, 2.5 mM MgCl_2_, 0.1 mg/mL BSA). In competitive binding experiments LANA peptide (1–23) was also added as a reaction component. Nucleosome was always added last. Prepared reactions were equilibrated on ice for 1 hr. Prior to sample loading, 6% TBE gels were run at 150 V for 60 min at 4°C in 0.25X Tris-borate-EDTA (TBE) buffer. Then, 10 μL of each equilibrated binding reaction was loaded on a 6% TBE gel and run at 150 V for 100-120 min at 4°C in 0.25X TBE buffer. Gels were stained in the dark on a rotating shaker for 20 min with SYBR Gold (Invitrogen) DNA-intercalating stain diluted to 1:5000, then imaged.

### Surface plasmon resonance

The affinity of RCC1 for nucleosome was measured on a Biacore 8K biosensor (GE Healthcare) by surface plasmon resonance (SPR). Streptavidin from Streptomyces avidinii (Millipore Sigma) corresponding to 2000 response units (RU) was amine-coupled on the utilized flow channels of a CM5 sensor chip. Approximately 200 RU of 5’ biotinylated 185 bp nucleosome (unmodified, H2AK119ub or H2BK120ub) was captured directly on flow cell 2 in separate channels. Binding experiments were carried out in HBS-EP+ buffer (10 mM HEPES pH 7.4, 150 mM NaCl, 3 mM EDTA, 0.05% Tween-20), where RCC1 was used as the analyte and titrated over flow cells 1 and 2 in two-fold dilutions. Following a 600 sec dissociation, an additional 300 sec injection of HBS-EP+ buffer served to regenerate the sensor surface. Sensorgrams were double-referenced against the control flow cell and buffer injections. Data were fit to a 1:1 steady-state affinity binding model using Biacore Insight Evaluation software and plotted in GraphPad Prism.

### Development of single domain antibodies (VHH) to the nucleosome acidic patch

Immunizations were performed at *Eurogentec*, phage display selection and recombinant expression at QVQ, and final characterization at *EpiCypher*. In brief, two llamas were immunized with nucleosomes and Freund’s Complete adjuvant, and boosted on days 28, 43, and 52 with additional immunogen and Freund’s incomplete adjuvant. RNA was harvested from peripheral blood lymphocytes on day 52, reverse transcribed, single domain antibody (also termed VHH; Variable Heavy Domain of a Heavy chain antibody) cDNAs isolated by PCR, and cloned to VCSM13 phage-display libraries of 2x10^9^ and 3x10^9^ transformants. After three rounds of library selection to nucleosomes, single-colony phage were isolated and ELISA screened to confirm target binding. Candidate VHH were Sanger sequenced and categorized to families. Chosen VHH (C-terminal 6His-tagged) were sequence verified, expressed as recombinants in BL21 *E.coli*, purified by Ni-NTA chromatography, and purity confirmed by SDS-PAGE. VHH epitopes were identified / characterized using fully defined nucleosomes (WT, PTM^+^, mutants) in dCypher Alpha (67) and dCypher Luminex (68).

### dCypher™ Luminex nucleosome binding assays

Semi-synthetic nucleosomes (Unmodified, Kub1-modified and acidic patch mutated) were separately coupled to saturation to distinct magnetic avidin-coated xMAP bead regions (*Luminex*). All handling and incubations of MagPlex beads were performed under subdued lighting. [Nucleosome: bead region] complexes were adjusted to 1 million beads / mL, multiplexed and exchanged into long term storage buffer (10 mM Cacodylate pH 7.5, 0.01% BSA, 0.01% Tween-20, 1 mM EDTA, 10 mM b-mercaptoethanol, 50% glycerol) for storage at -20°C. Panel balance and [Nucleosome : bead region] identity were confirmed using anti-dsDNA (EMD Millipore #MAB030; 1/5, 1/50 and 1/500), anti-H3.1/2 (Active Motif #61629; 1/250, 1/1000 and 1/4000) and anti-PTM (EMD Millipore #04-263, CST #8240S; both 1/250, 1/1000 and 1/4000).

For the dCypher Luminex assay, the binding of Queries (GST-2XLANA-His6 (hereafter GST-LANA) or VHH), to multiplexed nucleosomes (1000 beads/well: the Targets) was tested by serially diluting the Queries (two-fold: GST-LANA, 1.3 µM - 40 pM final; VHH, 2 µM - 2 nM final). In brief, 50 µL of multiplexed nucleosome panel was combined with 50 µL query in Luminex buffer (final: 20 mM Tris pH 7.5, 100 mM NaCl, 0.01% NP40, 0.01% BSA, 1 mM DTT) in a black, flat-bottom 96-well plate (*GreinerBio* #655900) and incubated under subdued lighting for one hr with shaking (800 rpm). Beads were captured / washed twice (100 µL of Luminex buffer) on a plate-based magnet. 100 µL of either anti-GST (*Fortis Life Sciences* #A190-122A; 1/1000) or Luminex buffer was respectively added to GST-LANA or VHH, and incubated for 30 min with shaking (800 rpm). Beads were captured / washed twice (100 µL of Luminex buffer) on a plate-based magnet. 100 µL of PE donkey anti-rabbit IgG (*BioLegend* #406421; 2 ug/mL) for GST-LANA, or PE goat anti-alpaca IgG VHH domain (*Jackson ImmunoResearch Laboratories* #128-115-232; 1/400) for VHH, was added and incubated for 30 min with shaking (800 rpm). After two additional washes, beads were resuspended in 100 µL Luminex buffer and Median Fluorescence Intensity (MFI) read on a FlexMap3D instrument (*PerkinElmer*), counting a minimum of 50-events per Target. Data was analyzed and visualized in *GraphPad Prism 10*.

The GST-LANA construct used was GST-TEVrs-2XLANA-2XGGGS-SrtAss-6HIS:

MSPILGYWKIKGLVQPTRLLLEYLEEKYEEHLYERDEGDKWRNKKFELGLEFPNLPYYIDGDVKLTQSMAIIRYIADKHNMLGGCPKERAEISMLEGAVLDIRYGVSRIAYSKDFETLKVDFLSKLPEMLKMFEDRLCHKTYLNGDHVTHPDFMLYDALDVVLYMDPMCLDAFPKLVCFKKRIEAIPQIDKYLKSSKYIAWPLQGWQATFGGGDHPPKSDLEVLFQGPLGSIEENLYFQSMAPPGMRLRSGRSTGAPLTRGSGGGGSMAPPGMRLRSGRSTGAPLTRGSGGGGSGGGGSLPATGGEGKSSGSGSESKSTGGHHHHHH

## RESULTS

### Ubiquitin conjugated to histone H2B-K120 occludes the nucleosome acidic patch

To investigate whether H2BK120ub adopts positions that could interfere with the binding of proteins that interact with the nucleosome acidic patch, we determined cryoEM structures of nucleosome core particles containing Widom 601 DNA (145 bp) and ubiquitinated histone H2B-K120 (Table 1). The density maps revealed four distinct positions of ubiquitin on the nucleosome surface, denoted H2Bub positions 1, 2, 3, and 4 (Figure 1A, Figure S1). In H2Bub position 1, ubiquitin sits nearest the center of the histone octamer core, and progressively further away in positions 2 – 4. The global resolution estimates for the EM maps for H2Bub positions 1, 2, 3, and 4 were calculated to be 3.32 Å, 3.33 Å, 3.34 Å, and 3.36 Å, respectively (Figures S2A-D). The local resolution of the nucleosome core, at ∼3 Å, was higher than that of the conjugated ubiquitin, at ∼6 Å, indicating greater conformational heterogeneity of the ubiquitin compared to the nucleosome core (Figures S3A-D). The secondary structure elements of ubiquitin in H2Bub positions 1-3 were sufficiently resolved to unambiguously position and orient the ubiquitin within the electron density maps (Figures S4A-C). In all three positions, the ubiquitin beta sheet is oriented towards the nucleosome surface, while the alpha helix faces away (Figure 1B). Given its proximity to the other positions, we modeled the ubiquitin in H2Bub position 4 in a similar orientation relative to the nucleosome surface (Figure S4D).

**Figure 1.**
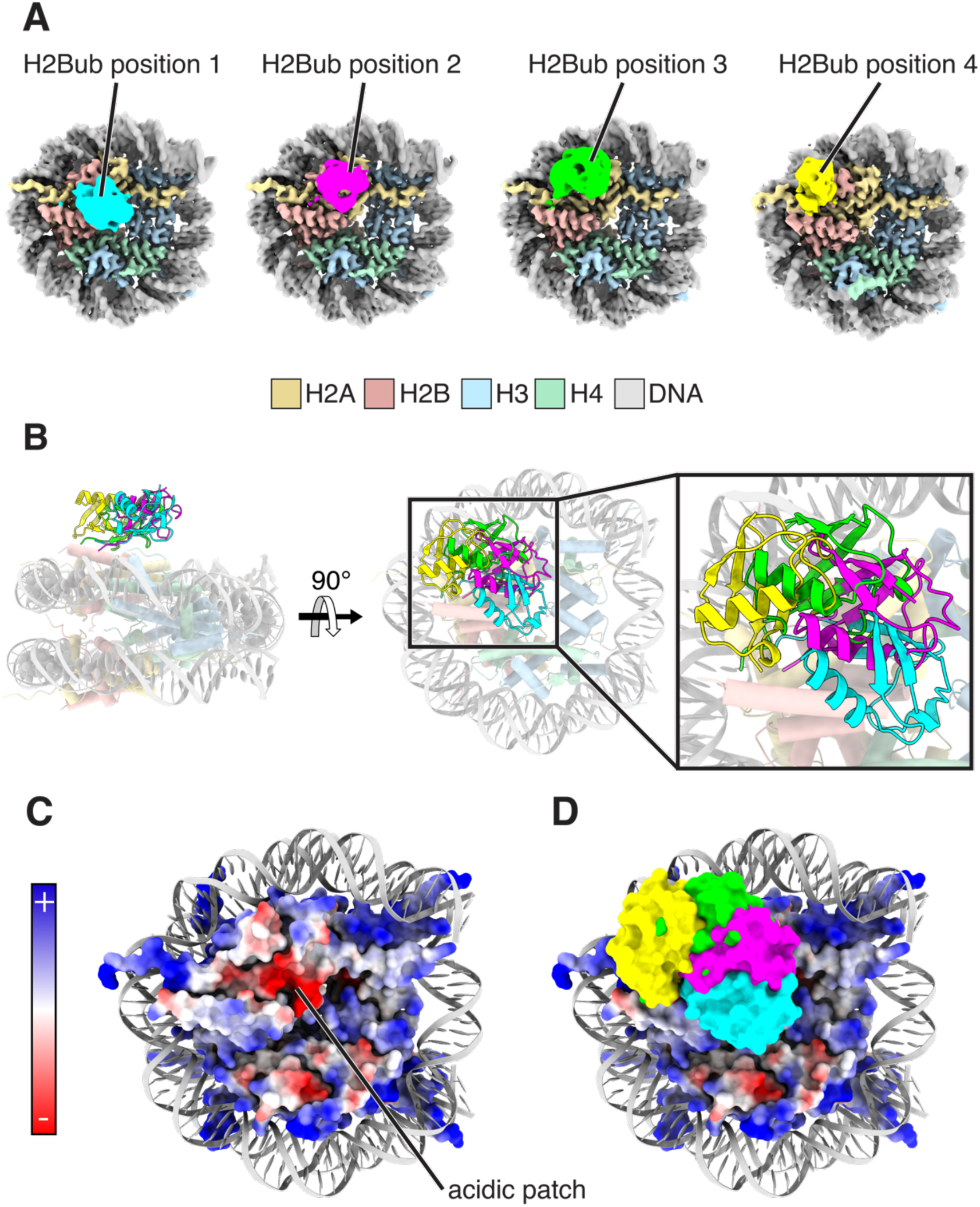
Four discrete ubiquitin positions in cryoEM maps of nucleosomes containing H2BK120ub. **(A)** CryoEM Maps of H2BK120ub nucleosome showing ubiquitin in four discrete positions. **(B)** Superposition of the four ubiquitin positions depicted in cartoon representation (colors as in panel A). **(C)** Electrostatic surface representation of the histone octamer core showing the nucleosome acidic patch. **(D)** Superposition of the four ubiquitin positions (colors as in A) over an electrostatic surface representation of the histone octamer core.

**Table 1.**
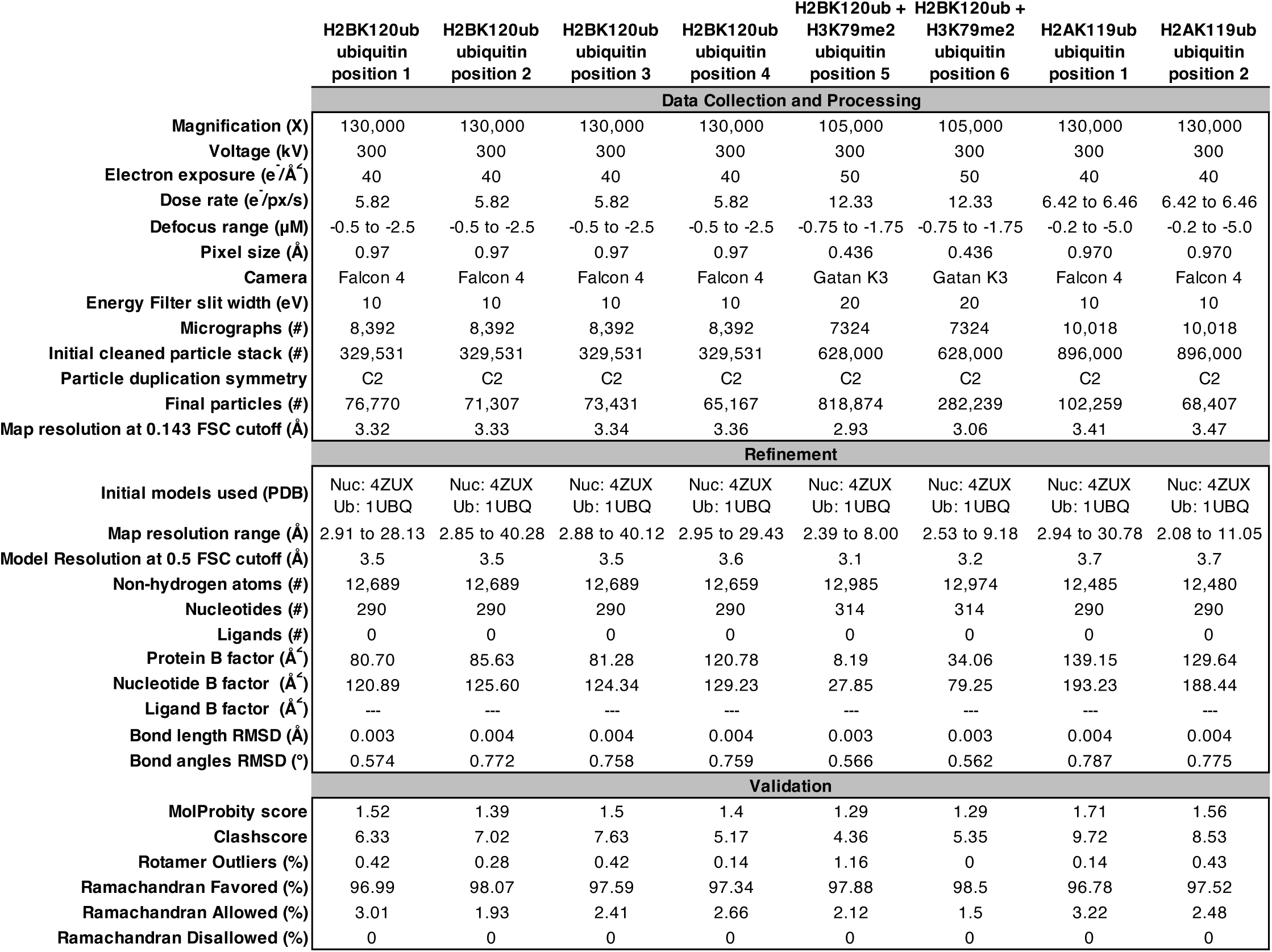
CryoEM data collection, processing, refinement, and validation statistics.

The ubiquitin in H2Bub positions 1 and 2 occludes large portions of the nucleosome acidic patch, near the center of the nucleosome face (Figure 1C). To quantitate the extent to which ubiquitin in each position occludes the acidic patch, we calculated the buried surface area between the acidic patch and ubiquitin using a range of probe radius values (see Methods) (Figures S5A-B). At a probe radius of 7 Å, the acidic patch surface area occluded by ubiquitin in H2Bub positions 1, 2, 3 and 4 is 1943 Å^2^, 1591 Å^2^, 1176 Å^2^, and 774 Å^2^, respectively. This trend is generally consistent across buried surface calculations performed using larger and smaller probe radii (Figures S5A-B).

Since methylation of histone H3K79 by DOT1L depends upon prior ubiquitination of histone H2BK120 (69), both modifications can co-occur on the same nucleosome (70). To see if this methyl modification affects H2BK120ub positioning on the nucleosome surface, we determined cryoEM structures of a nucleosome containing H2BK120ub, di-methylated histone H3-K79 (H3K_C_79me2), and Widom 601 DNA with 20 bp linkers (185 bp) (Table 1). The cryoEM dataset revealed two additional distinct ubiquitin positions on the nucleosome surface, denoted H2Bub positions 5 and 6 (Figures 2A-B, Figure S6). The global resolution estimates for these EM maps were 2.93 Å and 3.06 Å, respectively (Figures S2E-F), with a local resolution of ∼2.5 Å for the nucleosome core and ∼5 Å for ubiquitin (Figures S3E-F). The ubiquitin in position 5 lies between previously determined H2Bub positions 2 and 3 (Figure 2C). The ubiquitin in H2Bub position 6 lies closer to histone H3 than any of the other H2Bub positions, but does not interact directly with H3K_C_79me2. The density corresponding to ubiquitin in positions 5 and 6 was sufficiently well-resolved to unambiguously orient the ubiquitin with its beta sheet oriented towards the nucleosome surface (Figures 2A-B, Figures S4E-F). As observed for H2Bub positions 1 and 2, the ubiquitin in positions 5 and 6 occludes large portions of the nucleosome acidic patch (Figure 2D). At a probe radius of 7 Å, the occluded surface area between the acidic patch and ubiquitin in H2Bub positions 5 and 6 is 1599 Å^2^, and 1849 Å^2^, respectively (Figures S5A-B).

**Figure 2.**
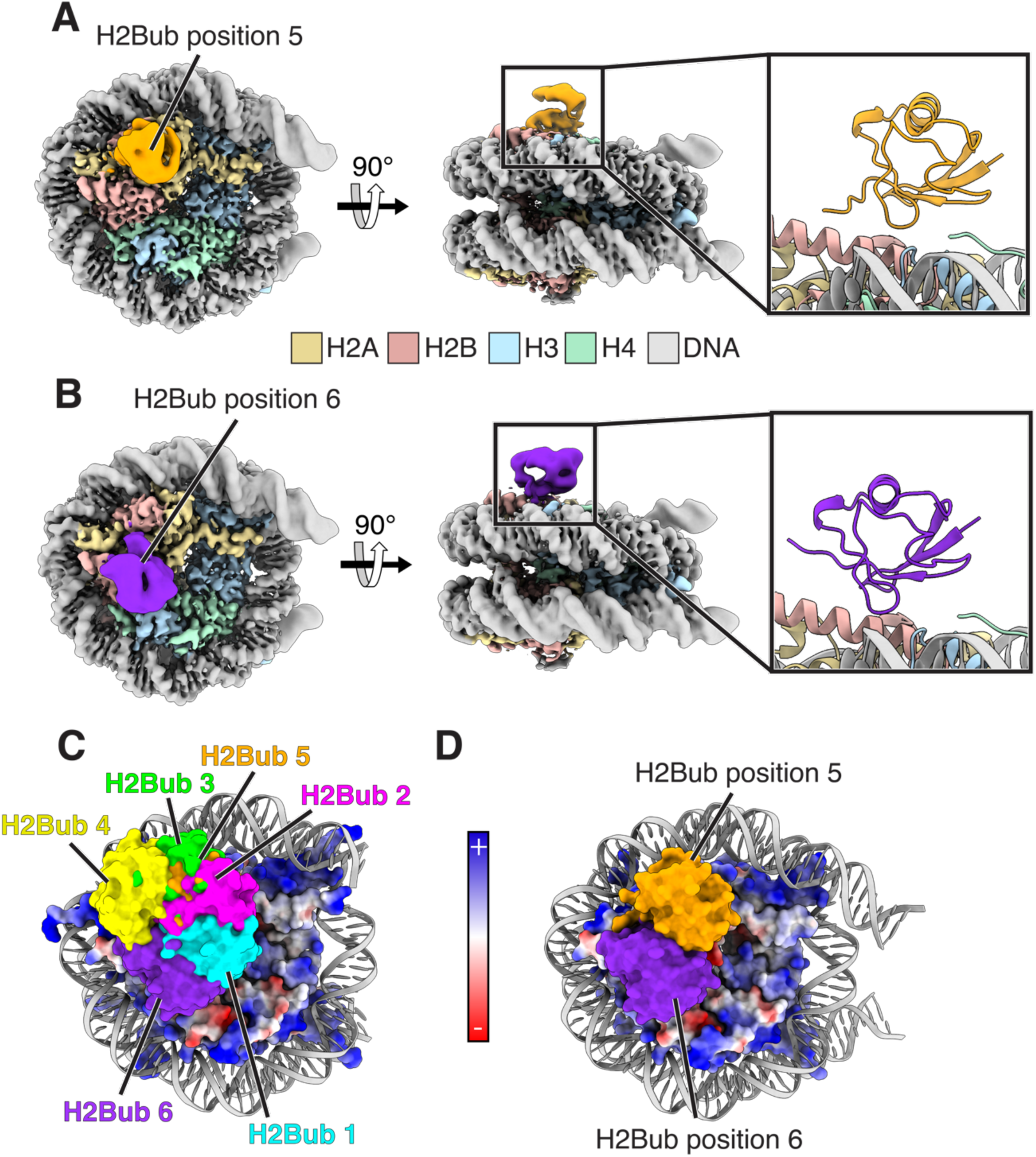
Two discrete ubiquitin positions in cryoEM maps of nucleosomes containing H2BK120ub and H3K_C_79me2. **(A)** CryoEM map of nucleosome containing H2BK120ub and H3K_C_79me2 with ubiquitin in position 5. Inset at right shows a cartoon model of H2Bub in position 5. **(B)** CryoEM map of nucleosome containing H2BK120ub and H3K_C_79me2 with ubiquitin in position 6. Inset at right shows a cartoon model of H2Bub in position 6. **(C)** The six ubiquitin positions observed in H2BK120ub nucleosomes with and without H3K_C_79me2 overlaid on a nucleosome with the histone octamer colored using an electrostatic surface representation. **(D)** Surface models of H2Bub in positions 5 and 6 overlaid on a nucleosome with the histone octamer colored using an electrostatic surface representation.

### Molecular dynamics simulations reveal stability of ubiquitin conjugated to H2B-K120

While our density maps indicate that ubiquitin conjugated to H2BK120 can adopt several discrete positions, these structural snapshots do not provide information on the dynamic motion of ubiquitin on the nucleosome surface. We therefore used molecular dynamics (MD) simulations to investigate the mobility of ubiquitin H2BK120ub-modified nucleosomes. We built the H2BK120ub-modified nucleosome system with the initial ubiquitin position at H2Bub position 5, which is approximately halfway between H2Bub positions 2 and 3. To gauge whether ubiquitin remained stably positioned over the acidic patch, we monitored the distance between their centers of mass over a 400 ns trajectory (Figure 3A). Over the course of the simulation, the distance between ubiquitin and the acidic patch varied between 24 Å and 36 Å (Figure 3B). Importantly, the ubiquitin primarily sampled conformations near the acidic patch and the distance between the ubiquitin and acidic patch largely remained within 1 Å of the starting ubiquitin position (Figure 3B). We note that ubiquitin samples some conformations farther from the acidic patch over the course of the simulation, as apparent from the local minima at ∼34 Å. This is consistent with our cryoEM data, where H2Bub can sample orientations with less acidic patch occlusion.

**Figure 3.**
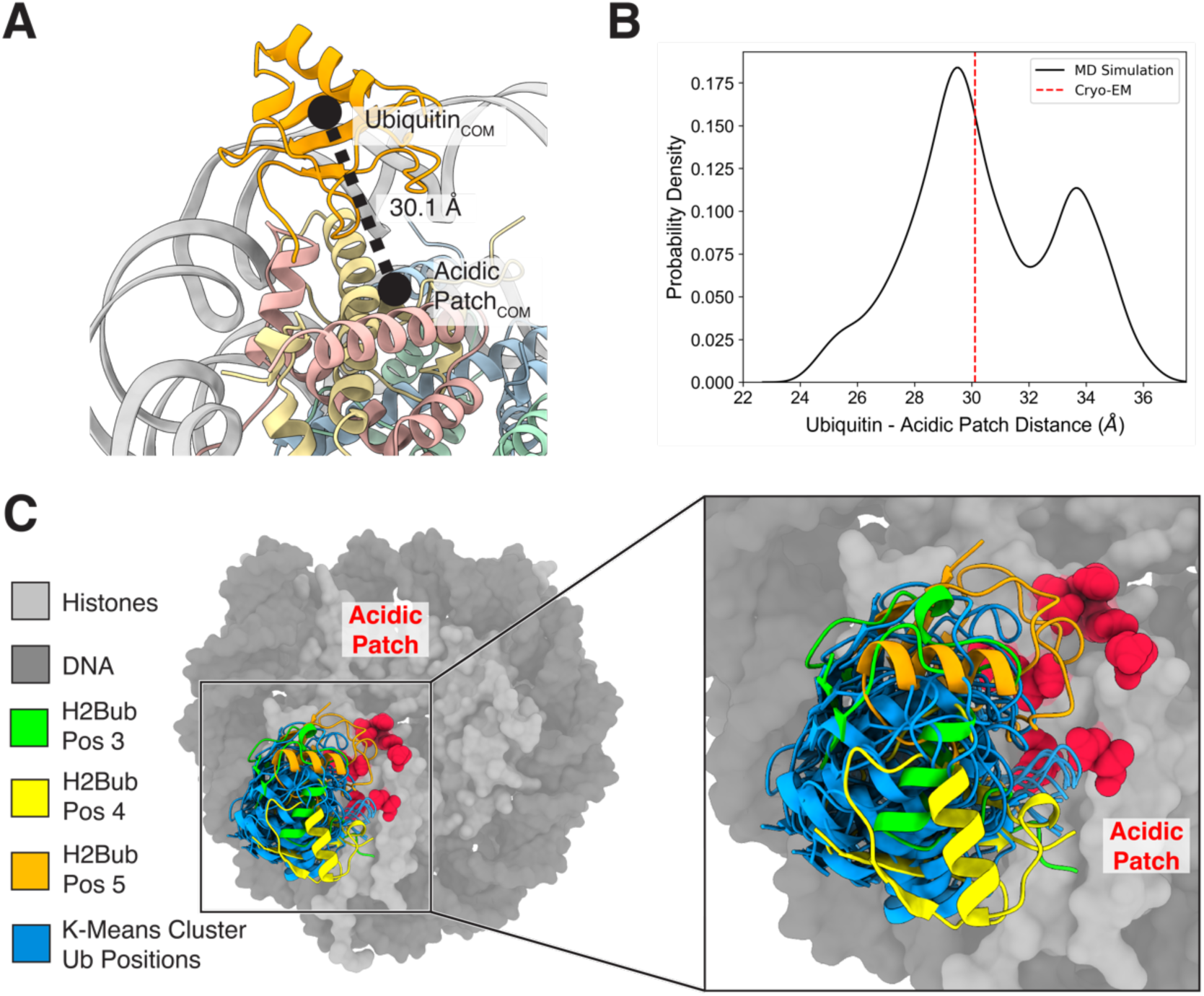
Molecular dynamics simulations H2BK120ub-modified nucleosome. **(A)** Diagram depicting calculation between the center of mass (COM) of ubiquitin and of the nucleosome acidic patch. The representative structure shows H2BK120ub nucleosome in the initial ubiquitin position for the MD simulations. **(B)** Probability density of finding ubiquitin at a specific distance from the acidic patch calculated over the entire simulation (black solid line). Initial starting position of ubiquitin indicated by a red dashed line. **(C)** K-means cluster analysis of ubiquitin showing the 10 most representative structures from one face of the nucleosome over the entire simulation (blue) and the ubiquitin cluster position relative to the nucleosome acidic patch (red), and ubiquitin in H2Bub positions 3 (green), 4 (yellow), and 5 (orange).

To visualize representative structures in the MD simulation, we used K-means clustering of ubiquitin throughout the trajectory to obtain representative snapshots consistent with H2Bub positions 3, 4, and 5 (Figure 3C). We did not see representative ubiquitin positions corresponding to H2Bub cryoEM positions 1, 2, and 6, likely because of limited sampling time. Overall, the MD simulations are consistent <u>with</u> our structural studies showing that ubiquitin conjugated to H2Bub occludes access to the acidic patch.

### Ubiquitin conjugated to histone H2A-K119 minimally occludes the nucleosome acidic patch

Ubiquitinated histone H2A-K119 (H2AK119ub) is one of the most abundant PTMs in higher eukaryotes (71), but its effect on protein binding to the nucleosome has not been explored. To see whether ubiquitin conjugated to H2A-K119 adopts discrete positions, we determined cryoEM structures of the nucleosome core particle containing ubiquitinated histone H2A-K119 and Widom 601 DNA (145 bp) (Table 1). The cryoEM dataset revealed two distinct ubiquitin positions, denoted H2Aub positions 1 and 2 (Figures 4A-B, Figure S7), with global resolution estimates for the corresponding EM maps of 3.41 Å and 3.47 Å, respectively (Figures S2G-H). Similar to the maps for H2BK120ub-modified nucleosome, the local resolution of the nucleosome core, at ∼3 Å, was higher than for conjugated ubiquitin, at ∼6 Å (Figures S3G-H). However, the map resolution was sufficient to unambiguously orient ubiquitin within the EM maps (Figures S4G-H).

**Figure 4.**
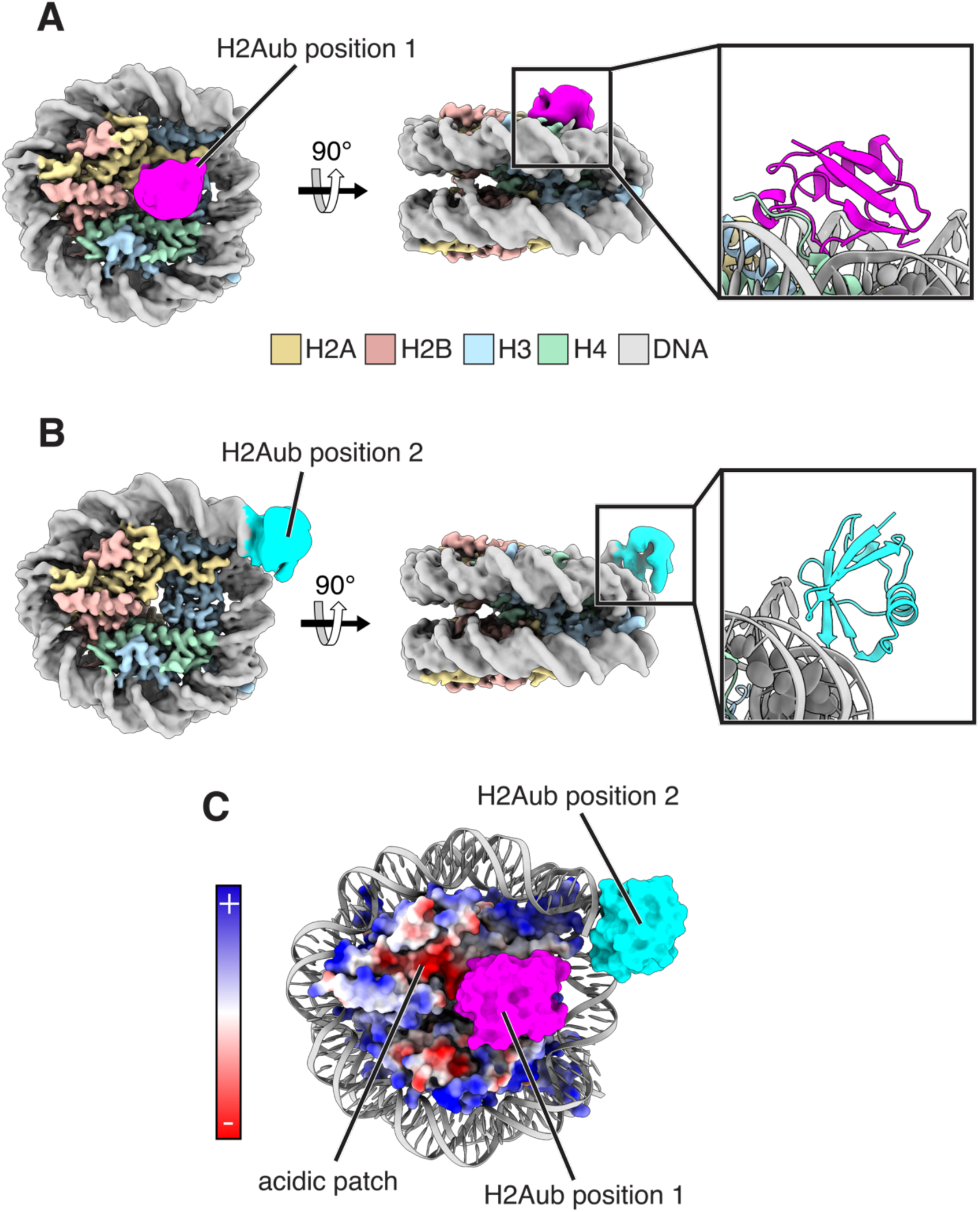
Two discrete ubiquitin positions in cryoEM maps of nucleosomes containing H2AK119ub. **(A)** CryoEM map of H2AK119ub nucleosome with ubiquitin in position 1. Inset shows a cartoon model of H2Aub in position 1. **(B)** CryoEM map of H2AK119ub nucleosome with ubiquitin in position 2. Inset shows a cartoon model of H2Aub in position 2. **(C)** CryoEM surface models of H2AK119ub nucleosome with both ubiquitin positions overlaid on a nucleosome with the histone octamer colored using an electrostatic surface representation.

The ubiquitin in H2Aub position 1 binds to the nucleosome surface over histones H3 and H4, immediately adjacent to the acidic patch (Figure 4C). In this position, ubiquitin partially overlaps the acidic patch, with an occluded surface area of 1241 Å^2^ as measured with a 7 Å probe radius (Figures S5A-B). The ubiquitin in H2Aub position 2 sits at the nucleosome dyad bound to the nucleosomal DNA terminus, end-on, with the ubiquitin beta-sheet oriented towards the DNA (Figure 4B). While the resolution is not sufficient to discern side chains, the ubiquitin position is consistent with van der Waals interactions between the hydrophobic patch on the ubiquitin beta sheet and the terminal base pair of the wrapping DNA. This ubiquitin position would be incompatible with a nucleosome embedded within a longer DNA duplex. We note that our attempts to determine the structure of H2AK119ub nucleosomes containing an additional 20 bp of flanking linker DNA (185 bp), which would be expected to prevent association of ubiquitin with the DNA end, were unsuccessful due to aggregation of the particles on cryoEM grids.

Since heterochromatic regions are characterized by chromatin that is condensed and enriched in H2AK119ub, (25) we explored whether the ubiquitin in H2Aub position 1 could be accommodated within higher-order nucleosome packing. While the precise nature of inter-nucleosomal interactions in condensed chromatin remains an area of active investigation (72), solution-based experiments, X-ray crystal structures, and cryoEM maps have revealed stacked nucleosomes (73–76). Modeling using the structure of a compacted tetranucleosome (73) shows that the ubiquitin in H2Aub position 1 can be readily accommodated between two stacked nucleosomes without steric clash (Figure S8). Ubiquitin in this position can also accommodate binding of the linker histone H1, which binds to the nucleosome dyad, promotes chromatin condensation and is abundant in heterochromatic regions (77,78).

### Ubiquitin conjugated to histone H2B but not H2A competes with RCC1 binding to nucleosomes

RCC1, a RAN GTPase (79), interacts with the nucleosome acidic patch and the adjacent DNA in a multivalent fashion (13). H2BK120ub has been shown to interfere with RCC1 binding to the nucleosome as assayed in pulldown experiments coupled with mass spectrometry analysis (32,33). It is therefore possible that H2BK120ub occlusion of the acidic patch observed in our structural studies might account for reduced nucleosome binding by RCC1. While the effect of H2AK119ub on RCC1 binding to nucleosomes has not been explored, we speculated this modification would have a minimal effect based on minimal occlusion of the acidic patch by H2AK119ub in our cryoEM structure (Figure 4).

We first quantitated the effect of H2BK120ub on RCC1 binding to unmodified and H2BK120ub nucleosomes using electrophoretic mobility shift assays (EMSA). On unmodified nucleosomes, increasing concentrations of RCC1 resulted in the formation of two discrete higher molecular weight bands (Figure 5A), which presumably correspond to 1:1 and 2:1 RCC1:nucleosome complexes. We confirmed the identify of these bands by showing that they disappear in the presence of increasing concentrations of LANA peptide, which binds to the acidic patch (35) and has been shown to abrogate RCC1 binding (80) (Figures S9A-B). At the highest concentrations of RCC1 (in the absence of LANA), we observed a high molecular weight smear resulting from non-specific aggregates containing RCC1 and nucleosome (Figure 5A). We next assayed RCC1 binding to H2BK120ub nucleosomes and found no discrete high molecular weight bands corresponding to RCC1:nucleosome complexes, although a high molecular weight smear corresponding to potential aggregates was visible at the highest RCC1 concentrations (Figure 5C). To rule out the possibility of artifacts due to the use of a non-native DCA linkage to attach ubiquitin to H2B-K120 (38), we confirmed that ubiquitin covalently linked a native isopeptide bond also disrupts formation of the 1:1 and 2:1 RCC1:nucleosome bands (Figure S9C).

**Figure 5.**
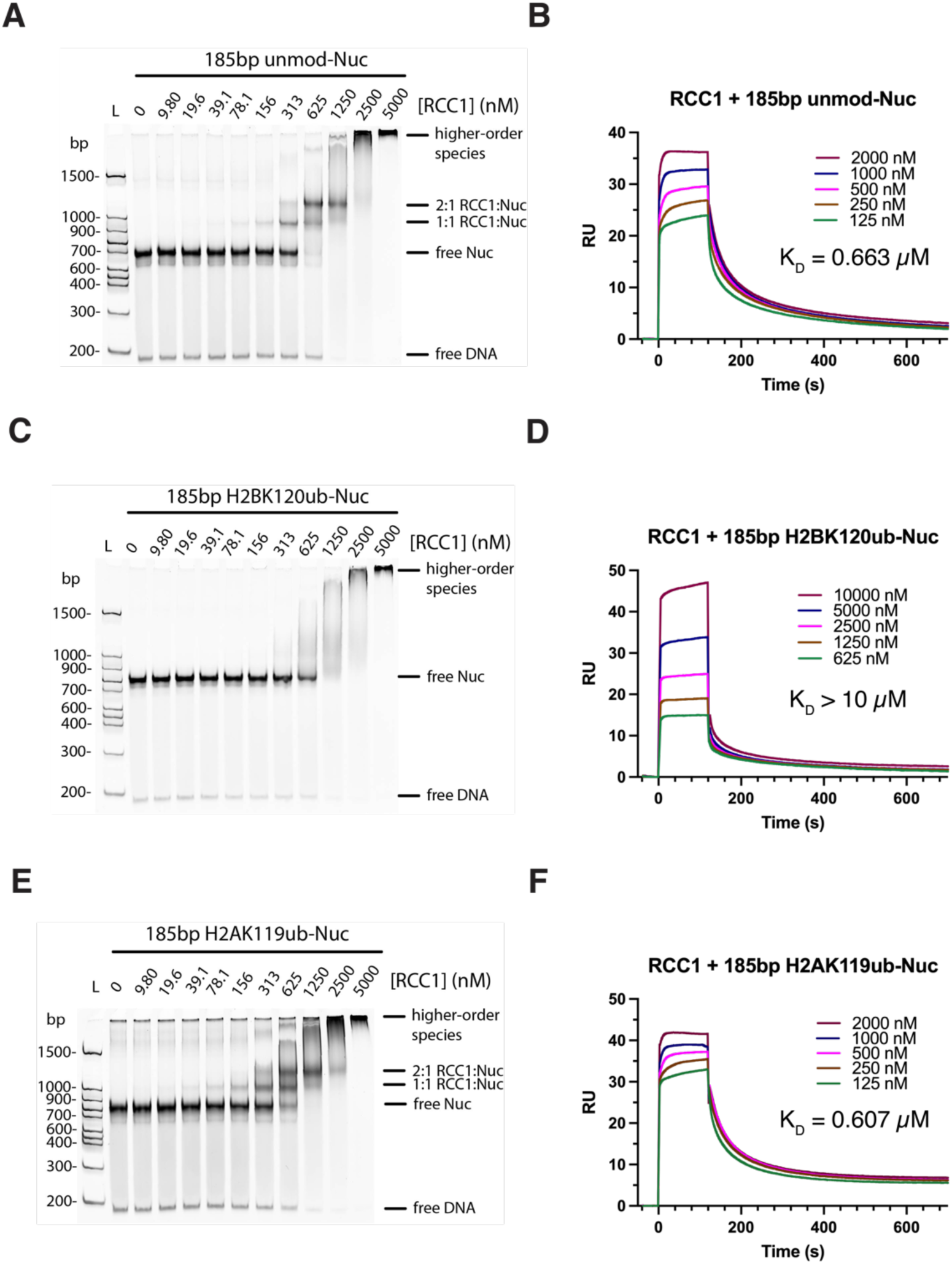
Binding of RCC1 to nucleosomes with ubiquitin conjugated to H2BK120 and H2AK119. **(A)** Electrophoretic mobility shift assay (EMSA) showing binding of the indicated concentrations of RCC1 to unmodified nucleosome (100 nM). Complexes visualized with DNA stain, SYBR Gold. **(B)** Surface plasmon resonance (SPR) assay of RCC1 binding to unmodified nucleosome. **(C)** EMSA showing binding of RCC1 to H2BK120ub nucleosome (100 nM). **(D)** SPR assay of RCC1 binding to H2BK120ub nucleosome. **(E)** EMSA showing binding of RCC1 to H2AK119ub nucleosome (100 nM). **(F)** SPR assay of RCC1 binding to H2AK119ub nucleosome.

To obtain a more precise measure of the impact of H2BK120ub on RCC1 binding to nucleosomes, we measured equilibrium dissociation constants by surface plasmon resonance (SPR). We found that RCC1 binds unmodified nucleosome with a K_D_ of 0.663 μM (Figure 5B), while the affinity for H2BK120ub nucleosomes is far lower, with an estimated K_D_ greater than 10 μM (Figure 5D). This far lower affinity is consistent with the reported role of H2BK120ub in interfering with RCC1 binding to nucleosomes (32,33).

To test whether it is simply the presence of ubiquitin that interferes with binding to nucleosomes, we assayed RCC1 binding to H2AK119ub nucleosomes by EMSA. Increasing the concentration of RCC1 results in formation of discrete bands corresponding to 1:1 and 2:1 RCC1:H2AK119ub nucleosome complexes (Figure 5E), similar to those observed in assays of binding to unmodified nucleosome (Figure 5A). The K_D_ of RCC1 for nucleosomes containing H2AK119ub was measured by SPR to be 0.607 μM (Figure 5F), similar to the K_D_ of RCC1 for unmodified nucleosomes (Figure 5B). These results indicate that ubiquitin conjugated to H2AK119 does not interfere with RCC1 binding to the nucleosome, consistent with our cryoEM data showing that ubiquitin in H2AK119ub nucleosomes minimally occludes the nucleosome acidic patch (Figure 4C).

To further explore the ability of H2BK120ub or H2AK119ub to interfere with nucleosome binding, we compared the effect of these PTMs on additional acidic patch-binding entities. Nucleosome binding by a glutathione-S-transferase-LANA fusion protein (GST-LANA) decreases in the presence of H2BK120ub (34), while the impact of H2AK119ub has not been explored. We used dCypher™ Luminex (see Methods) to compare the binding of GST-LANA to an unmodified, H2BK120ub, H2AK119ub, and acidic patch mutant H2A(E61A) nucleosome panel in multiplex. As expected, the binding affinity of GST-LANA was significantly weaker on H2BK120ub or H2A(E61A) nucleosomes than on unmodified nucleosomes (Figure 6A, Table 2). As observed for RCC1 (Figure 5E), GST-LANA binding to H2AK119ub nucleosomes was indistinguishable from unmodified (Figure 6E).

**Figure 6.**
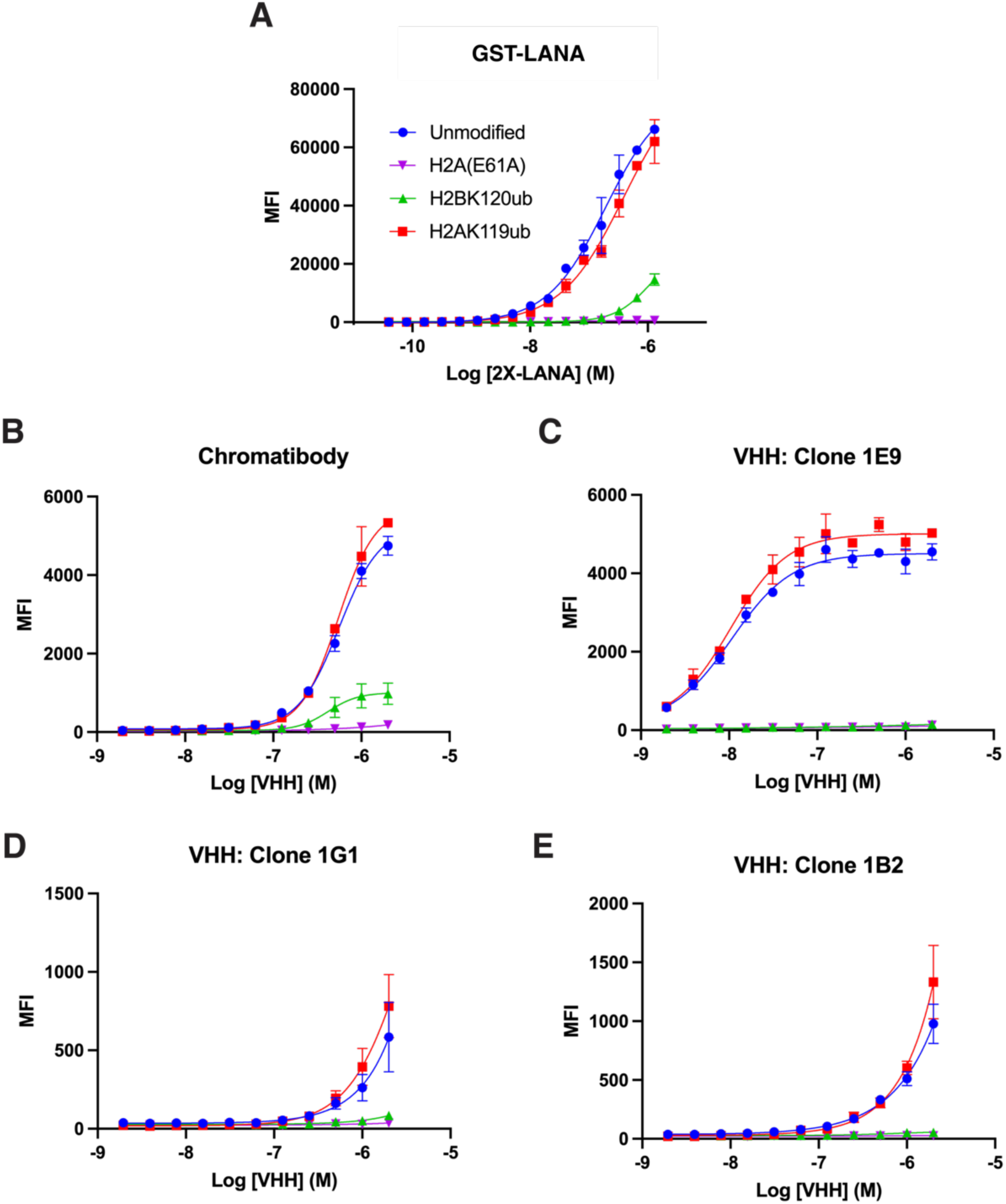
H2BK120ub but not H2AK119ub inhibits interactions with the nucleosome acidic patch. Acidic patch mutation H2A(E61A) and H2BK120ub, but not H2AK119ub, interferes with the binding of GST-LANA **(A)** (34) and chromatibody VHH **(B)** to nucleosome (95) in dCypher-Luminex assays (see Methods); measured using median fluorescence intensity (MFI) units. **C-E)** H2A(E61A) and H2BK120ub, but not H2AK119ub, also interfere with the binding of three newly generated nucleosome acidic patch specific VHH (clones 1E9, 1G1 and 1B2: see Methods).

**Table 2.** EC50 values in dCypher-Luminex binding assays.

|  | EC50 (nM) |  |  |  |
| --- | --- | --- | --- | --- |
|  | Unmodified | H2A(E61A) | H2BK120ub | H2AK119ub |
| **GST-LANA** | 192 | NB | > 1300 | 392 |
| **Chromatibody** | 559 | NB | 419 | 544 |
| **1E9** | 11 | NB | NB | 10 |
| **1G1** | > 1000 | NB | NB | > 1000 |
| **1B2** | > 1000 | NB | NB | > 1000 |

We next assayed the effect of H2BK120ub or H2AK119ub on the binding of four single domain antibodies (aka. Variable Heavy Domain of a Heavy chain antibodies; VHH) that specifically bind the nucleosome acidic patch (see Methods). As shown in Figures 6B-E, the H2A(E61A) acidic patch substitution abrogated nucleosome binding by all four VHH (chromatibody, 1E9, 1G1, 1B2), confirming the importance of acidic patch contacts. As observed for RCC1 and GST-LANA, the presence of H2BK120ub substantially reduced VHH-nucleosome binding, while binding to H2AK119ub nucleosomes was indistinguishable from unmodified nucleosomes (Figures 6B-E). These results indicate that H2BK120ub, but not H2AK119ub, generally interferes with nucleosome engagement by acidic patch binding entities.

## DISCUSSION

Our results provide a mechanism by which ubiquitin conjugated to histone H2B-K120 can exclude proteins from the nucleosome. There had been reports that nucleosome binding by human RCC1, yeast SIR3, and the viral LANA peptide was disrupted by monoubiquitination of histone H2BK120 (32–34). These proteins bind the nucleosome acidic patch (13,35,36), leading to the speculation that H2BK120ub may act as a gatekeeper to this cluster of acidic residues in histones H2A/H2B (32,34,81,82). Our cryoEM maps unexpectedly showed ubiquitin positioned on the nucleosome in several distinct locations that either partly or completely occlude the nucleosome acidic patch (Figure 1, Figure S5). The resulting steric clash could explain how H2BK120ub interferes with binding of select proteins to the nucleosome acidic patch (2). By contrast, cryoEM maps of H2AK119ub nucleosomes revealed two stable ubiquitin positions, only one of which slightly overlapped the acidic patch. Figures 5 and 6). The consistent effect of H2B-K120 ubiquitination on nucleosome binding by unrelated proteins is consistent with the idea that any gatekeeping function of H2BK120ub is steric, and not related to particular folds or sequence motifs.

It is not clear why the ubiquitin conjugated to H2B-K120 adopts discrete positions over the acidic patch in our EM maps. In reported structures of proteins that bind to both the histone octamer and the ubiquitin in H2BK120ub, ubiquitin adopts different positions that depend, in each case, on direct interactions with the bound protein (83–87). A crystal structure of a nucleosome containing H2BK120ub alone did not show density for ubiquitin (88), indicating either that ubiquitin is conformationally heterogeneous in the absence of a bound protein, or that it adopts multiple positions that could not be resolved in the electron density maps. Interestingly, a previous cryoEM structure of the Chd1 chromatin remodeler bound to a nucleosome containing H2BK120ub and partially unwrapped DNA shows EM density for ubiquitin over the acidic patch (81). However, it was not clear if the partially unwrapped DNA or the presence of Chd1 contributed to ubiquitin positioning, although we note that the density maps do not show direct interactions between the remodeler and ubiquitin.

It is important to note that the relative residence time of ubiquitin at each observed position cannot be determined from cryoEM maps. While it is most likely that ubiquitin coupled to H2B-K120 has many degrees of freedom and can adopt a large number of positions relative to the nucleosome, our ability to capture discrete snapshots at sufficient resolution to orient the ubiquitin suggests it has sufficient residence time at or near the acidic patch to interfere with protein binding. Molecular dynamics simulations of H2BK120ub nucleosomes support our observation that ubiquitin largely remains near the acidic patch (Figures 3A-B). Although our MD simulations were able to recapitulate H2Bub positions 3, 4, and 5 observed in our cryoEM maps, we did not capture positions 1, 2, and 6, suggesting that further sampling is required to observe them. Future work using longer simulations or enhanced techniques will be needed to recapitulate other H2Bub positions, and potentially identify metastable states that cannot be captured by cryoEM. Nevertheless, our MD simulations interrogate the dynamics of H2BK120ub and show that the ubiquitin remains positioned at or near the nucleosome acidic patch, thereby occluding this interaction hub.

Although the precise nature of the contacts between ubiquitin and the nucleosome could not be resolved, the EM maps for nucleosomes containing H2BK120ub position 1 and 2 show weak density for ubiquitin residue R74 interacting with nucleosome acidic patch residues H2A-E56 and H2B-E113 (Figure S10). It is possible these interactions help position the ubiquitin on the nucleosome, although we note that there is no density for R74 interaction in density maps corresponding to H2Bub positions 3 through 6. The ubiquitin hydrophobic patch, composed of beta sheet residues L8, I44, and V70 (89), is a common site of interaction with ubiquitin binding proteins (89–91). In our cryoEM structures of H2BK120ub nucleosomes, the ubiquitin hydrophobic patch is oriented towards the nucleosome surface (Figures 1B and 2A-B). Further investigations will be needed to determine which ubiquitin residues are most important for H2BK120ub positioning.

Importantly, we observed stable positioning of ubiquitin on H2BK120ub and H2AK119ub nucleosomes in the absence of additional proteins. DOT1L, Dot1, and COMPASS all bind to ubiquitin conjugated at H2BK120, as well as to the nucleosome acidic patch (34,83,84,92–94). The ubiquitin is thus repositioned as a result of energetically favorable interactions with each protein, clearing their way to contact the nucleosome acidic patch. Such multivalent binding to both ubiquitin and the nucleosome could explain why DOT1L, Dot1, and COMPASS, which contain ubiquitin binding elements, bind specifically to nucleosomes containing H2BK120ub, while RCC1, Sir3, and LANA, which do not bind ubiquitin, are excluded from the nucleosome acidic patch by this PTM.

Monoubiquitination of histones H2A and H2B play opposing roles in regulating transcription through their interaction with enzymes that specifically recognize these modifications, and by promoting or disrupting chromatin compaction (15,16). Our studies shed light on a potential additional mechanism in which H2BK120ub, but not H2AK119ub, regulates binding to the acidic patch, a hub of nucleosome engagement for diverse chromatin interactors.

## DATA AVAILABILITY

Models and cryoEM maps were deposited in the Protein Data Bank (PDB) and Electron Microscopy Data Bank (EMDB) under the following accession codes: H2BK120ub-modified nucleosome ubiquitin position 1 (PDB: 8V25, EMDB: 42898), position 2 (PDB: 8V26, EMDB: 42899), position 3 (PDB: 8V27, EMDB: 42900), position 4 (PDB: 8V28, EMDB: 42901); H2BK120ub+H3K79me2-modified nucleosome ubiquitin position 5 (PDB: 8G6G, EMDB: 29767), position 6 (PDB: 8G6H, EMDB: 29769); H2AK119ub-modified nucleosome ubiquitin position 1 (PDB: 8G6Q, EMDB: 29778), position 2 (PDB: 8G6S, EMDB: 29781).

Raw cryoEM movies will be deposited in the EMPIAR database.

## SUPPLEMENTARY DATA

Supplementary Data are available at NAR online.

## AUTHOR CONTRIBUTIONS

Chad W. Hicks: Conceptualization, Formal analysis, Investigation, Methodology, Resources, Validation, Visualization, Writing—original draft. Sanim Rahman: Formal analysis, Investigation, Methodology, Validation, Visualization, Writing—review & editing. Susan L. Gloor: Formal analysis, Investigation, Methodology, Validation, Visualization. James K. Fields: Investigation, Formal analysis, Methodology, Validation. Natalia Ledo Husby: Resources, Validation. Anup Vaidya: Resources, Validation. Keith E. Maier: Resources, Validation. Michael Morgan: Resources. Michael-Christopher Keogh: Project administration, Funding acquisition, Supervision, Writing – review & editing. Cynthia Wolberger: Project administration, Funding acquisition, Supervision, Writing—review & editing.

## ACKNOWLEDGEMENTS

We thank Duncan Sousa and Dazhong (David) Ding for advice and support with cryoEM sample preparation and data collection at the Beckman Center for Cryo-EM at Johns Hopkins. We thank Edvin Pozharskiy for advice and support with cryoEM grid screening at the Maryland Center for Advanced Molecular Analysis (M-CAMA). We thank the Wolberger lab for their insights and discussions on the manuscript.

---

**Figure S1.**
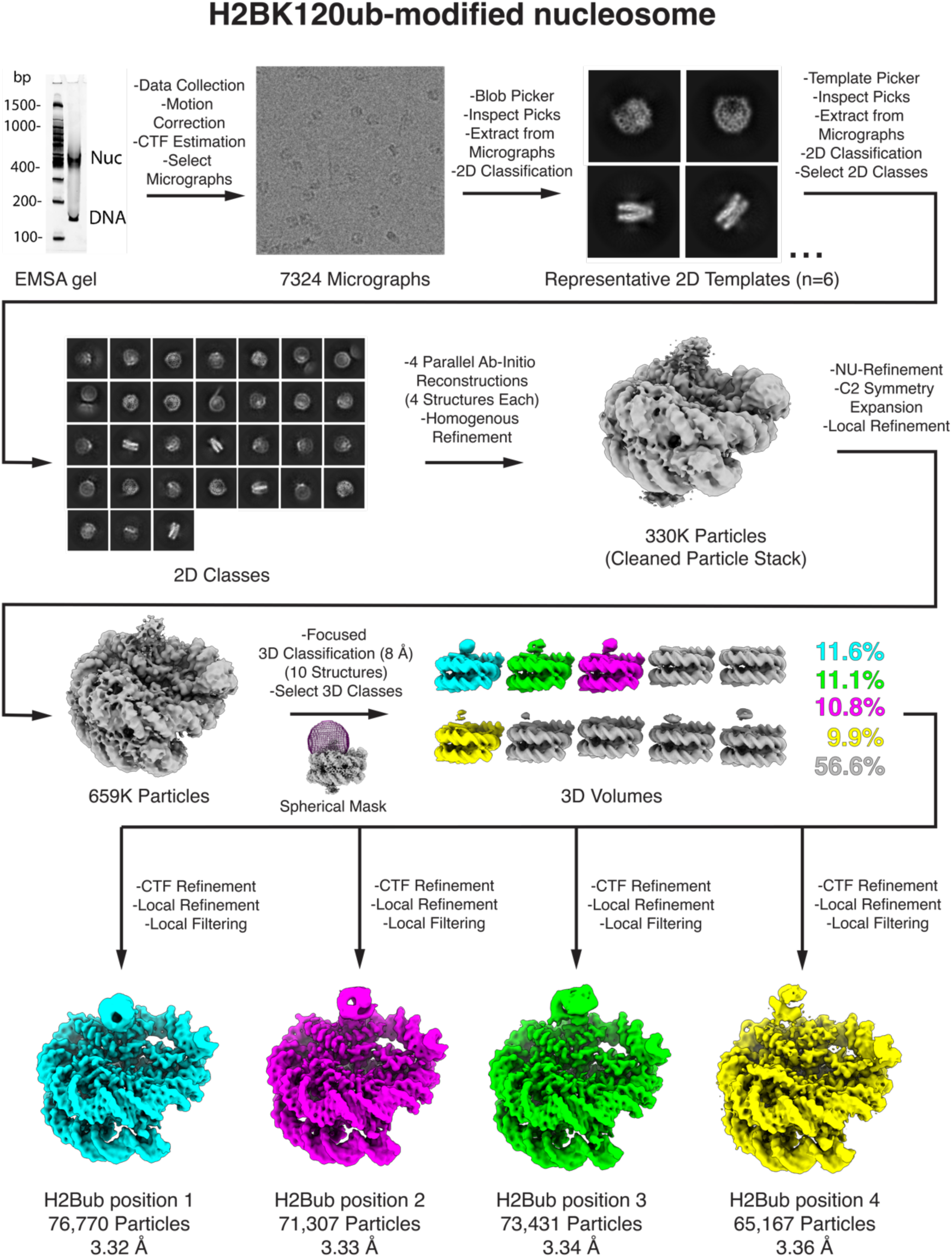
CryoEM data processing workflow for nucleosome containing H2BK120ub and Widom 601 DNA (145 bp).

**Figure S2.**
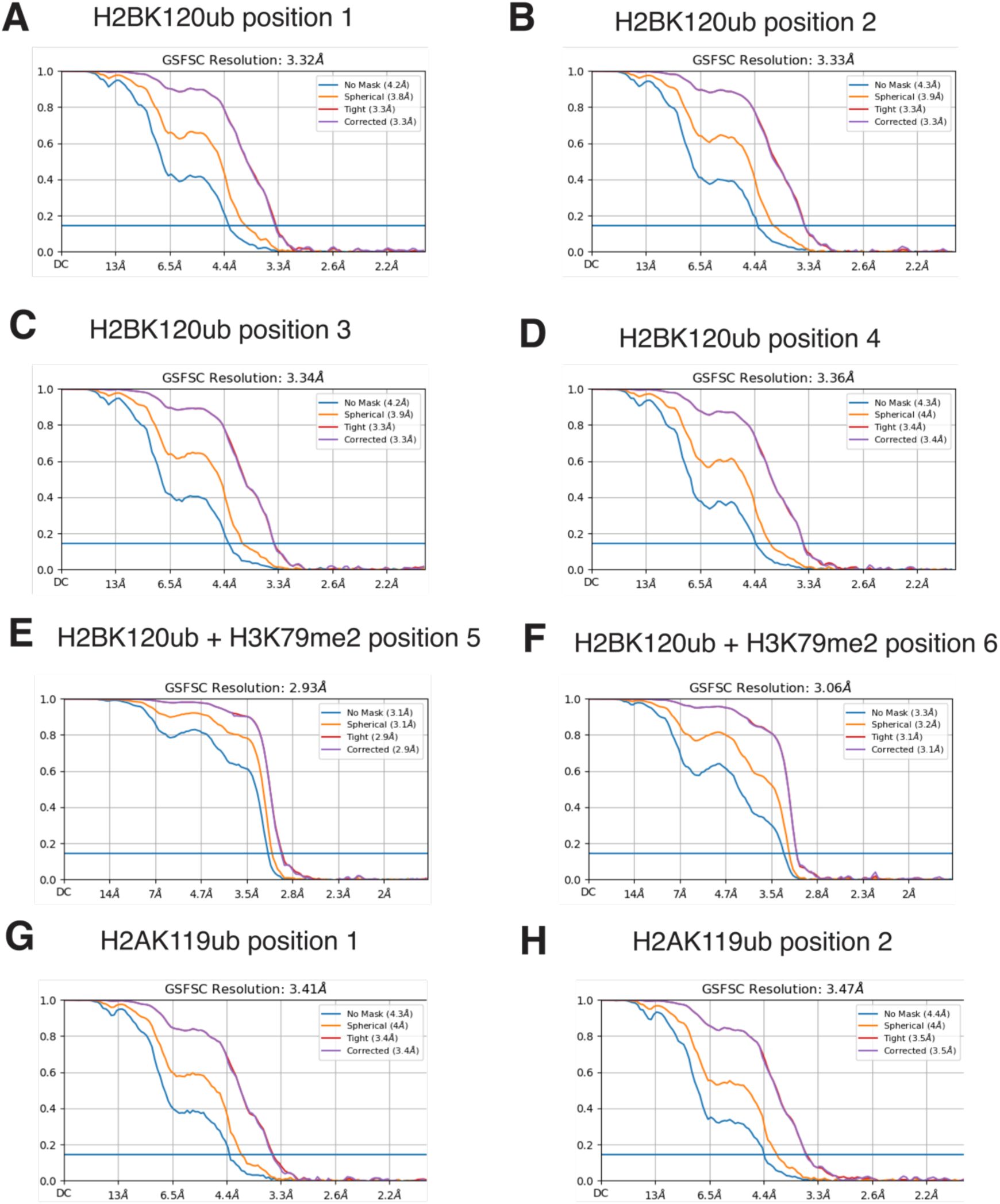
Fourier Shell Correlation (FSC) plots of all cryoEM maps. FSC plots using gold-standard 0.143 cutoff of cryoEM structures for: H2BK120ub nucleosome **(A-D)**, H2BK120ub+H3K_C_79me2 nucleosome **(E-F)**, and H2AK119ub nucleosome **(G-H)**.

**Figure S3.**
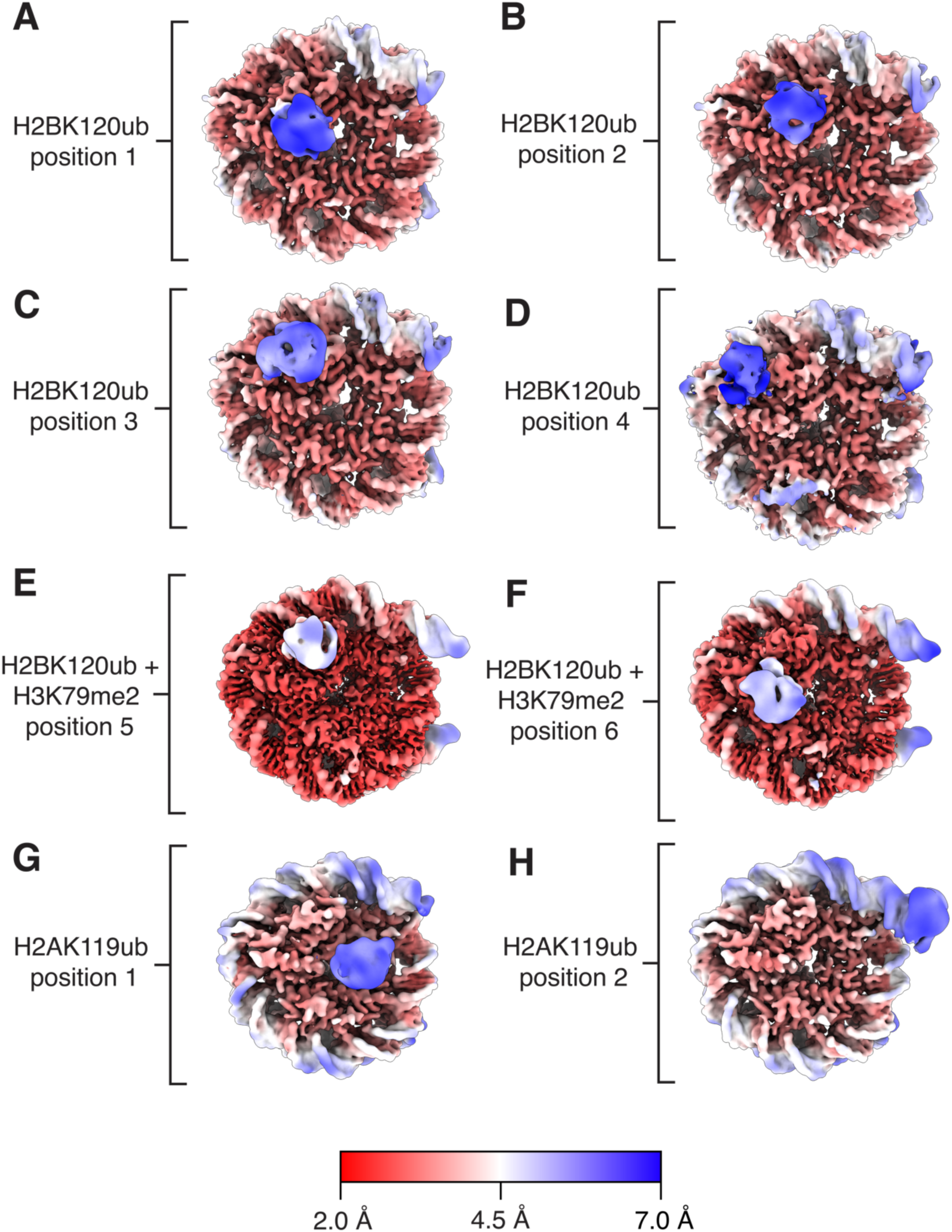
Local resolution estimation representations of all cryoEM maps. Local resolution estimation color depictions of cryoEM maps of H2BK120ub nucleosome **(A-D)**, H2BK120ub+H3K_C_79me2 nucleosome **(E-F)**, and H2AK119ub nucleosome **(G-H)**.

**Figure S4.**
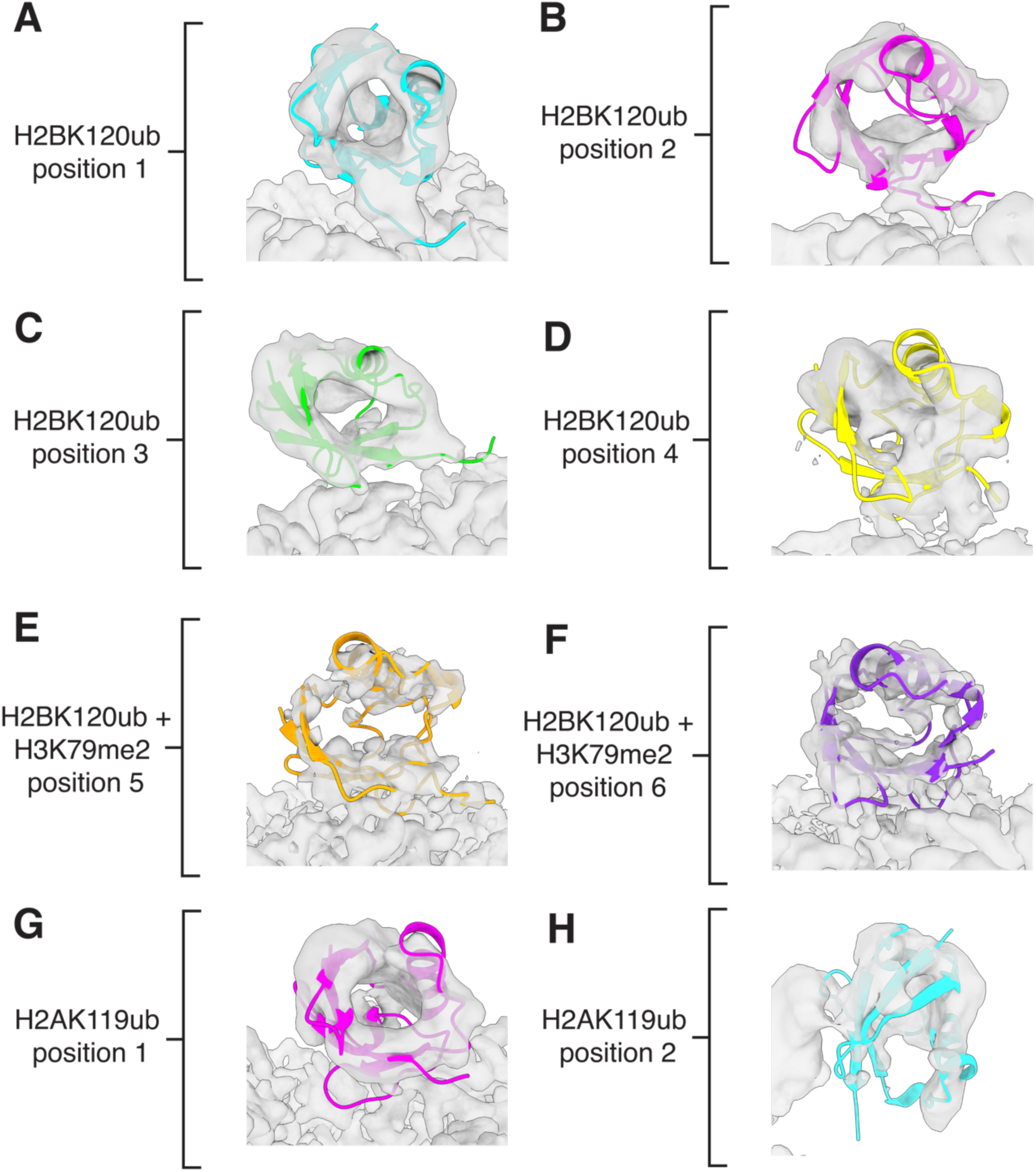
Fit of model to map for all structures. Ubiquitin cartoon model showing the fit to cryoEM maps corresponding to: H2BK120ub nucleosome, with ubiquitin in positions 1, 2, 3, and 4 **(A-D)**, H2BK120ub+H3K_C_79me2 nucleosome with ubiquitin in positions 5 and 6 **(E-F)**, and H2AK119ub nucleosome with ubiquitin in positions 1 and 2 **(G-H)**.

**Figure S5.**
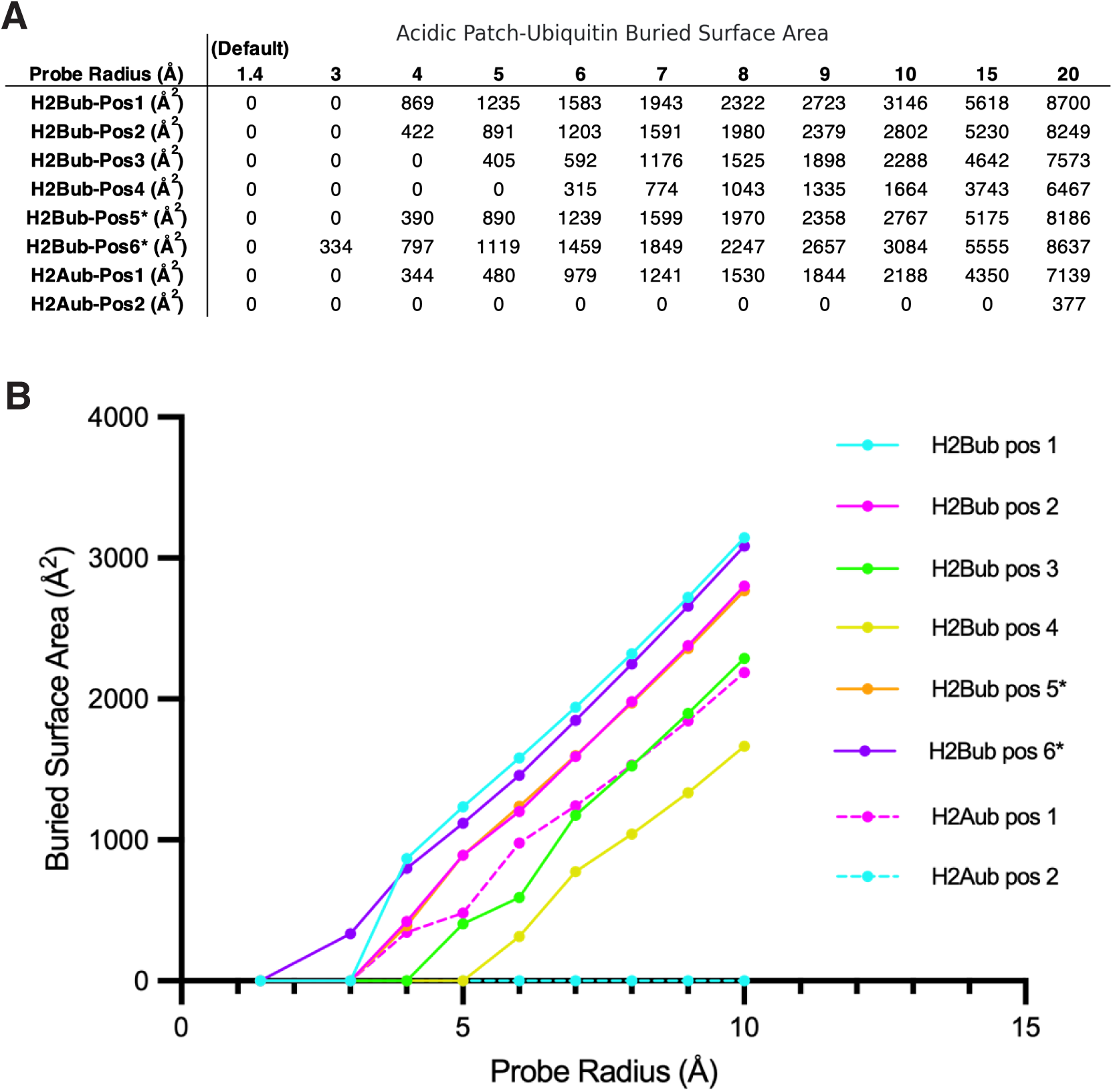
Acidic patch – ubiquitin buried surface area calculations for all ubiquitin positions. Full table of values **(A)** and line chart (up to 10 Å probe radius) **(B)** showing buried surface area between ubiquitin (all Ub residues) and the nucleosome acidic patch (H2A residues E56, E61, E64, D90, E91, E 92 and H2B residues E105, E113) at all ubiquitin positions. Calculated using “interfaces” command in ChimeraX at a range of probe radii. Solid lines correspond to H2Bub positions while dashed lines and square data points correspond to H2Aub positions. * indicates the H2Bub position was derived from the H2BK120ub+H3K_C_79me2 nucleosome cryoEM dataset.

**Figure S6.**
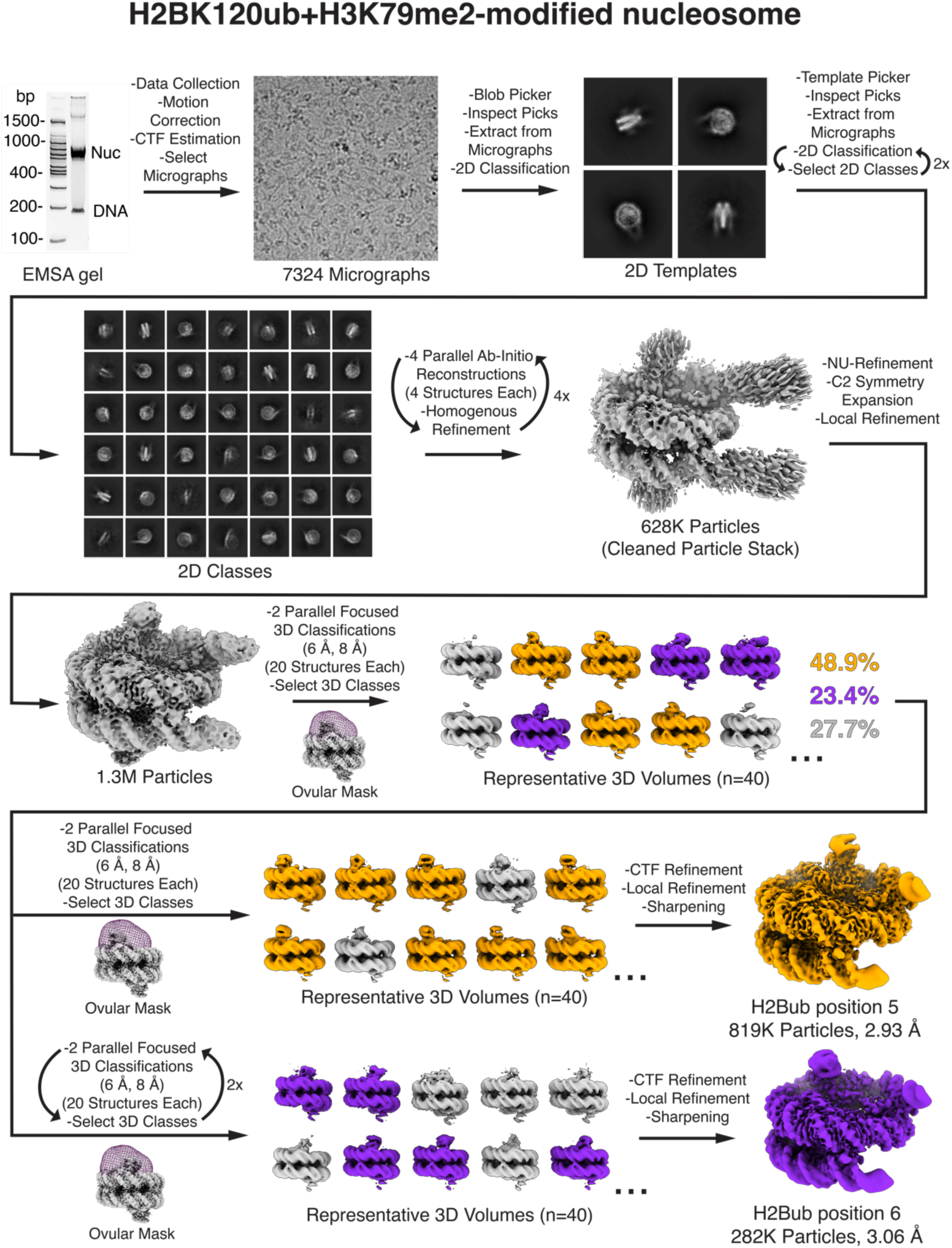
CryoEM data processing workflow for nucleosome containing H2BK120ub, H3K_C_79me2, and Widom 601 DNA with 20 bp linkers (185 bp).

**Figure S7.**
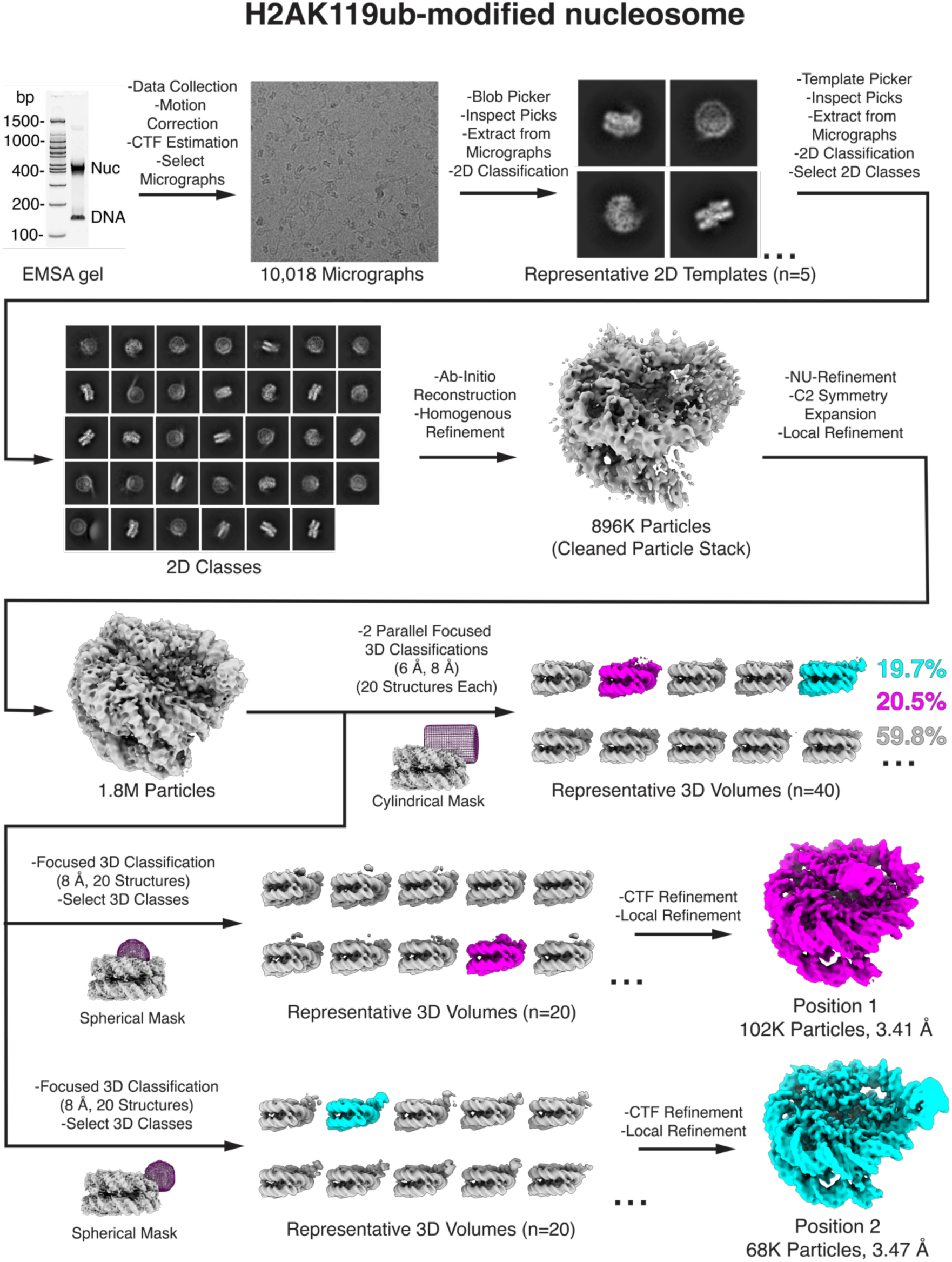
CryoEM data processing workflow for nucleosome containing H2AK119ub and Widom 601 DNA (145 bp).

**Figure S8.**
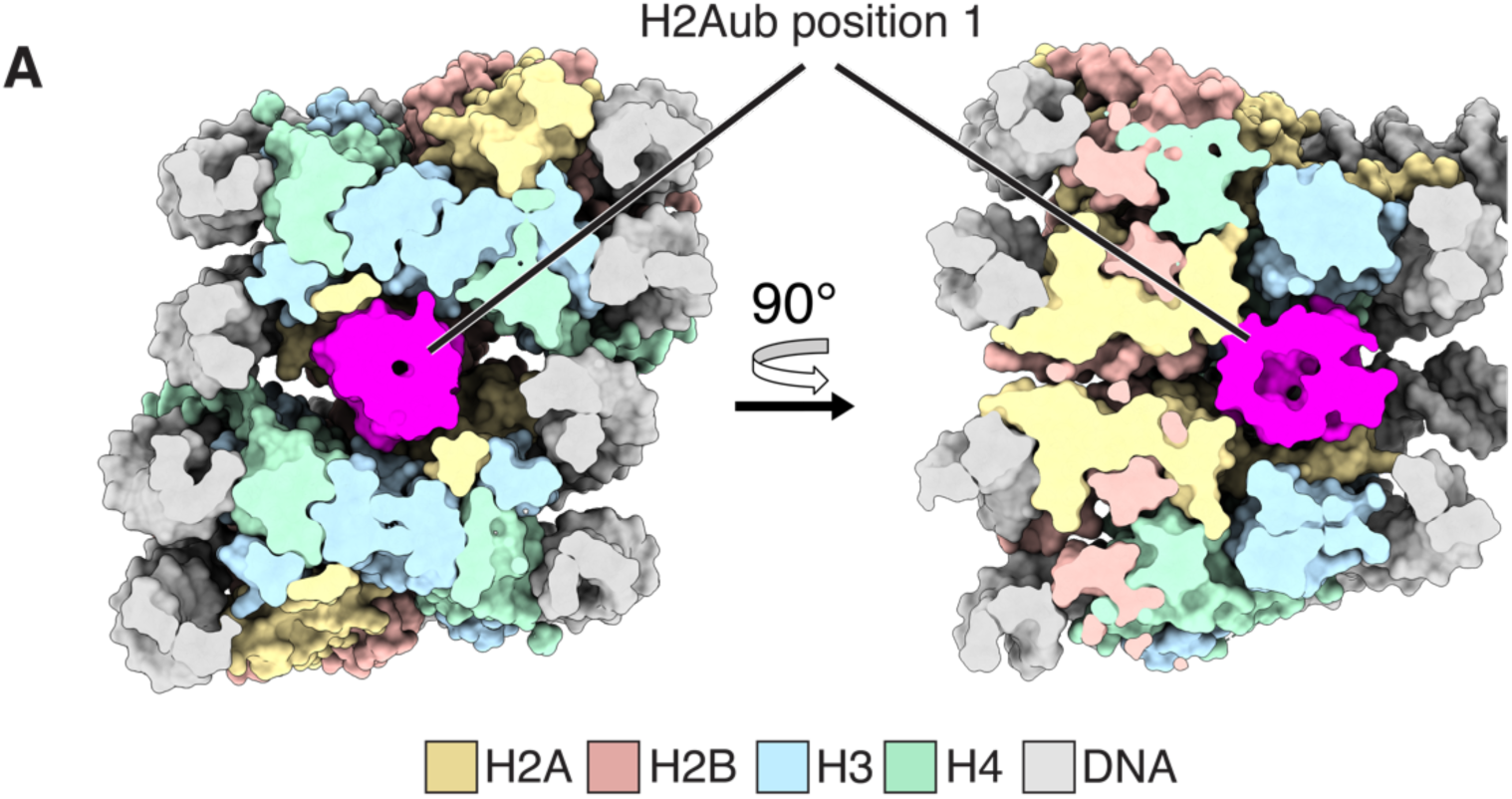
H2AK119ub nucleosome with ubiquitin in position 1 can accommodate nucleosome stacking without steric clash. CryoEM model of the ubiquitin of H2Aub nucleosome in position 1 (ubiquitin + lower nucleosome), depicted in surface representation, superimposed over an X-ray structure of a tetranucleosome (PDB:1ZBB). Figure is shown in a cut-away surface representation, with the slice halfway through the ubiquitin.

**Figure S9.**
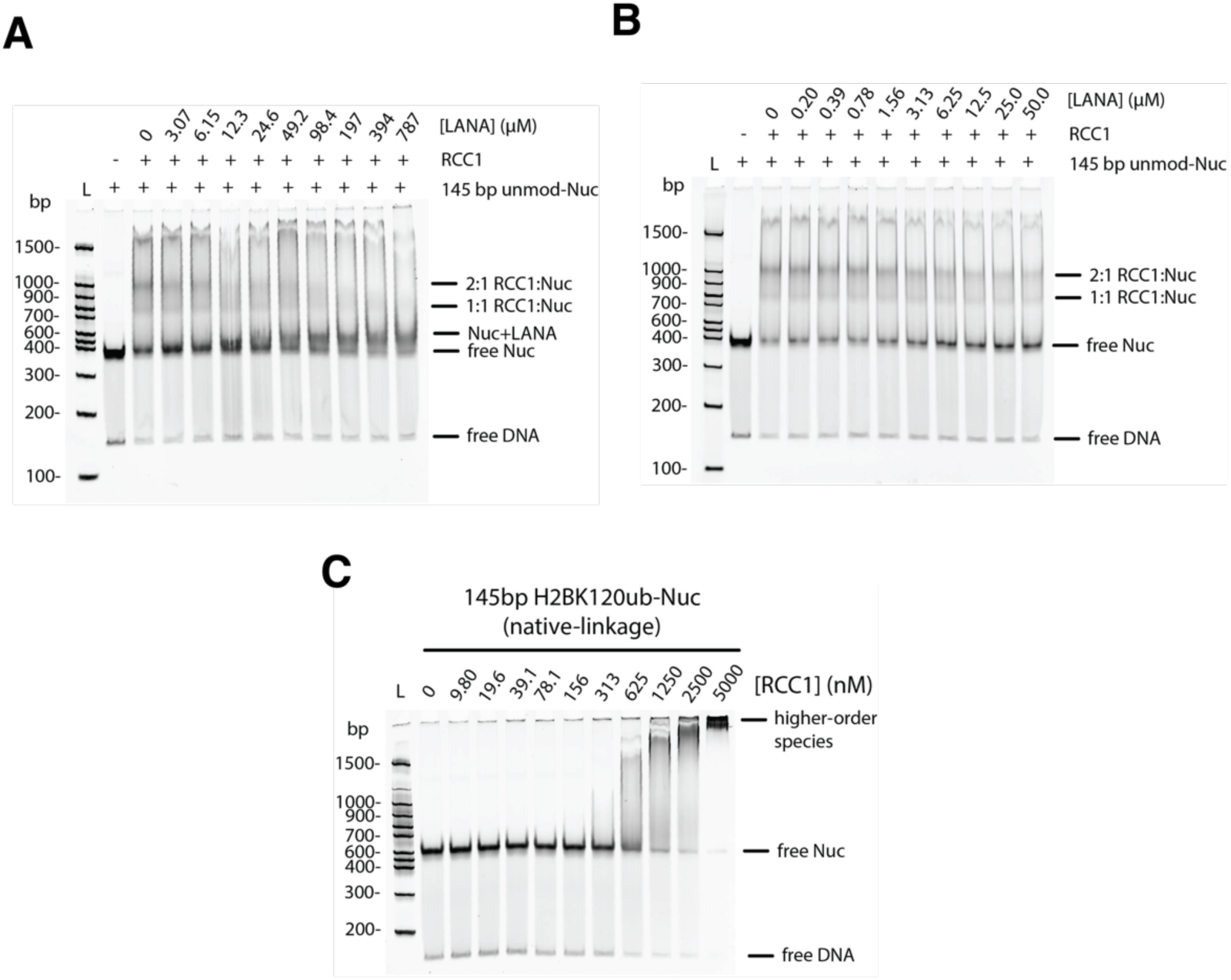
Binding of RCC1 to nucleosomes in the presence and absence of LANA peptide. **(A)** EMSA showing RCC1 (625 nM) binding to unmodified nucleosome (100 nM) in the presence of increasing amount of LANA peptide (0-787 mM). **(B)** As in (A), with 0-50 mM LANA peptide. **(C)** RCC1 binding to nucleosomes containing a native linked H2BK120Ub. The nucleosomes in all panels contained 145 bp Widom 601 DNA. All gels stained with SYBR Gold.

**Figure S10.**
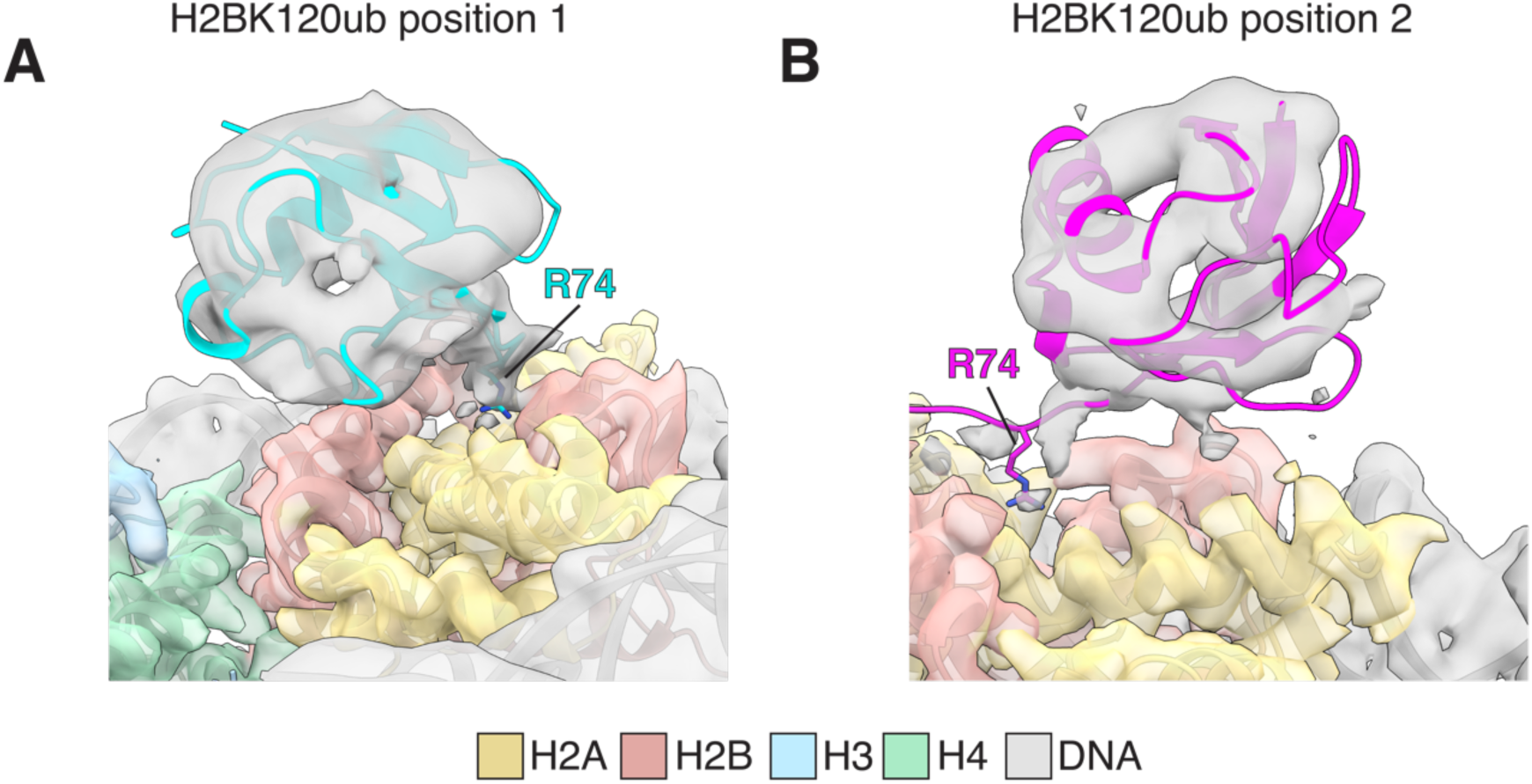
Arginine 74 of ubiquitin in H2BK120ub nucleosome interacts with the acidic patch formed by histone H2A/H2B. **(A)** R74 of ubiquitin in position 1 interacts with the nucleosome surface. **(B)** R74 of ubiquitin in position 2 interacts with the nucleosome surface.

